# Cyfip2 mediates sensorimotor integration of visual input through Rac1-dependent actin remodeling

**DOI:** 10.64898/2025.12.21.695802

**Authors:** Kimberly M. Charron, D. Christopher Cole, Melody B. Hancock, Sureni H. Sumathipala, Audrey L.B. Johnson, Katya J. Frazier, Kurt C. Marsden

**Affiliations:** North Carolina State University, Biological Sciences, Raleigh, NC; California State University San Marcos, San Marcos, CA

**Keywords:** *cyfip2*, zebrafish, behavior, visual system, development, cytoskeletal remodeling

## Abstract

Sensorimotor integration of visual input is essential for adaptive behaviors such as navigation, hunting, and escape from danger, yet our understanding of the developmental mechanisms that assemble these visuomotor circuits remains incomplete. *Cytoplasmic FMRP-interacting protein 2 (cyfip2)* is a key factor in retinotectal axon guidance in zebrafish larvae, and it has conserved roles in regulating actin dynamics and mRNA translation. Variants in human *CYFIP2* cause neurodevelopmental disabilities including vision deficits, but how it functions to shape visually-driven behavior is unclear. Here we measured multiple visually-mediated behaviors, brain-wide activity, and conditional rescue of behavioral defects in larval zebrafish to define *cyfip2*’s roles in assembling visual sensorimotor circuits. *cyfip2* mutants display severe deficits in prey capture and dark and light flash responses, despite normal optokinetic responses, indicating intact retinal phototransduction but disrupted downstream sensorimotor integration. Both spontaneous and stimulus-evoked neuronal activity are reduced in the optic tectum of *cyfip2* mutants, as measured by phospho-ERK immunostaining and pan-neuronal calcium, consistent with impaired functional input from retinal ganglion cells. Finally, using conditional transgenic alleles to temporally control Cyfip2 expression, we identified a critical window from 30-50 hours post-fertilization during which Cyfip2 acts through Rac1-dependent actin polymerization but not FMRP-mediated translation to establish visual behavior circuits. Together, these findings define how and when Cyfip2 acts as a critical organizer of vertebrate visual circuit development, providing mechanistic insight into the visual and motor deficits in human subjects with variants in CYFIP2.

## Introduction

Sensorimotor integration forms the foundation for essential survival skills, from rapid escape responses to complex navigation, by transforming streams of sensory input into precisely timed and spatially coordinated motor output. In vertebrates, the optic tectum (orthologous to the mammalian superior colliculus) serves as a hub for visuomotor processing by integrating retinotopically mapped visual inputs from retinal ganglion cells (RGCs) with contextual cues, and through outputs to hindbrain motor circuits the tectum drives context-specific behavioral responses that underlie perception and action^1–3^. Assembly of these neural circuits requires a coordinated set of cellular and molecular processes to drive proper neuronal specification and migration, axon growth and guidance, and synapse formation and function to enable a wide range of visually-driven adaptive behaviors.

To construct an accurate map of the visual field in the brain, axonal projections from RGCs to their targets in the optic tectum must be topographically ordered, and this organization is controlled by a complex genetic program of attractive and repulsive cues and cytoskeletal regulators^4–7^. One of these genetic factors is *cytoplasmic Fragile X Messenger Ribonucleoprotein Interacting Protein 2 (cyfip2)*, which was identified through a forward genetic screen for RGC axon pathfinding defects in larval zebrafish^8–11^. In wild-type fish, dorsonasal RGCs axons are sorted into the ventral branch of the optic tract and enter the tectum ventrally where they terminate and arborize, but in *cyfip2* mutants, a subset of dorsonasal axons are missorted into the dorsal branch of the optic tract and incorrectly enter the tectum dorsally^8,11,12^. These missorted axons take circuitous routes through the lateral tectum but appear to eventually terminate and arborize correctly in the ventral tectum^8,11^.

*cyfip2* is a key regulator of both actin cytoskeletal remodeling and mRNA translational control, two processes critical for axon guidance and synapse formation. As a member of the WASP-family verprolin-homologous protein (WAVE) heteropentameric regulatory complex (WRC)^13–17^, *cyfip2* binds to activated GTP-bound Rac1, releasing WASF from the WRC to initiate Arp2/3-dependent branched actin polymerization, driving neuronal morphogenesis, axon guidance, and growth cone dynamics^15,18,19^. Through interactions with binding partners Fragile X Messenger Ribonucleoprotein (FMRP) and eukaryotic translation initiation factor 4E (eIF4E), *cyfip2* also controls local translation of neuronal mRNAs, including of transcripts important for learning and memory and synapse formation^20,21^. In the visual system, *cyfip2* regulates dorsoventral RGC axon sorting and guidance into the tectum through WRC mediated actin dynamics, independently of FMRP-dependent translational regulation^12^. How these mechanisms and the relatively subtle defects in RGC axon pathfinding impact the function of visual circuits is unknown, however.

Here, we directly test whether *cyfip2* is required for visually-driven behavior and interrogate the relationship between the genetic control of axon pathfinding and behavioral outcomes by linking molecular pathways, critical developmental windows, and neural activity with functional consequences for visual sensorimotor integration. Our findings demonstrate that, during a critical developmental window, *cyfip2*-dependent actin remodeling is required for the maturation and function of sensorimotor circuits that enable visually guided behavioral responses in larval zebrafish.

## Materials & Methods

### Ethics statement

All zebrafish experiments were carried out ethically and with the approval and oversight of North Carolina State University’s IACUC (protocol #23-368). The North Carolina State University IACUC follows the US Public Health Service’s Policy on Humane Care and Use of Laboratory Animals and Guide for the Care and Use of Laboratory Animals.

### Zebrafish husbandry and maintenance

Adult zebrafish were maintained in 5 fish/L density under a light/dark cycle of 14:10 h at 28°C, with a recirculating water system kept at pH of 7.0-7.3, and a conductivity of 420-460 μS. Animals were fed GEMMA Micro 300 (Skretting) in the morning, followed by *artemia* brine shrimp (Brine Shrimp Direct) in the afternoon. The *cyfip2^p4^*^00^ line was maintained from a forward genetic screen^22^, the heatshock-inducible transgenic lines *Tg(hsp70:cyfip2-GFP)^p407tg^*, *Tg(hsp70:cyfip2-(C179R)-GFP)^ncu102Tg^*, and *Tg(hsp70:cyfip2-(K723E)-GFP)^ncu103Tg^* were maintained following creation as previously described^22,23^. All fish lines were maintained by outcrossing to Zebrafish International Resource Center (ZIRC)-derived AB wildtype fish.

Zebrafish larvae were generated from pairwise crosses in mating boxes (Aquaneering) containing system water and artificial grass. Embryos were collected the following morning, 2-3 hours into the light cycle. For time-sensitive experiments, pairs were separated by a gate until the start of the light cycle at 8am, when the gate was released and embryos were collected after one hour. All embryos were collected into petri dishes containing 1X E3 media (5 mM NaCl, 0.17 mM KCl, 0.33 mM CaCl_2_ · H_2_0, 0.33 mM MgSO_4_), then sorted under a brightfield microscope for fertilization and normal development at ∼6 hpf into dishes of ≤ 65 embryos per 100 mm Petri dish. Embryos were raised on a light/dark cycle of 14:10 h at 29°C, 50% of 1x E3 media was changed daily, and any embryos with gross morphological defects were removed and euthanized. All experimental data was obtained from larvae prior to sexual differentiation.

### DNA extraction and genotyping

Following experimentation, larvae were individually fixed with 100% methanol. Following methanol evaporation, DNA was extracted from whole larvae using the HotShot DNA lysis method consisting of a lysis buffer (25 mM NaOH, 0.2 mM EDTA) followed by a neutralization solution (40 mM Tris-HCl). Genotyping for *cyfip2^p4^*^00^, *GFP, cyfip2^C179R^, and cyfip2^K723E^* was performed by PCR amplification using primers described in Table 5-1 followed by gel electrophoresis or Sanger Sequencing.

### Inducible Heatshock Rescue

To induce expression of *Tg(hsp:cyfip2-GFP)*, 20 larvae were incubated in 1.5 mL of E3 media per Eppendorf tube, and placed in a thermal water bath at 38°C for 40 min at either 12, 30, 54, 78, 96, or 120 hpf. For continuous expression of *Tg(hsp:cyfip2-GFP)* larvae were repeatedly placed in a thermal water bath at 38°C for 40 min at 30,54,78, and then 102 hpf. To determine molecular mechanisms of action, expression of *Tg(hsp:cyfip2-GFP)* was induced by incubating larvae in 38°C thermal bath for 25 min at 30 hpf to provide GFP fluorescence comparable to the C179R and K723E variants after 40 min at 38°C. Following heatshock, larvae were returned to petri dishes and allowed to recover until behavioral testing at 5 dpf.

To compare induced protein expression between *Tg(hsp:cyfip2-GFP)* and the C179R and K723E variants, 20 larvae were incubated in 1.5 mL of E3 media per Eppendorf tube at 12 hpf, and placed in a thermal water bath at 38°C for a duration of 25,40, 60, or 80 minutes, and returned to petri dishes with pronase and allowed to recover for 5 hours. Larval fluorescence expression was then imaged 5 hours post-heatshock with a Nikon SMZ25 stereo microscope with a GFP bandpass filter, Lumen 200 illumination system, and Nikon DS-Qi2 monochrome microscope camera. Image analysis was performed using FIJI to manually create ROIs of the larval body (excluding the yolk sac and eye) and eye. Fluorescence intensity values are reflective of mean gray values recorded for respective ROIs.

### Brain activity and morphometry analysis

To capture whole-brain baseline activity and morphology of free-swimming larvae, phosphorylated-ERK antibody staining was conducted as previously described^24,25^. Larvae were fixed in 4% paraformaldehyde (PFA) diluted into 1x Phosphate Buffered Saline (PBS). Following overnight fixation, larvae were washed with 1X PBS with 0.5% Triton (PBST), bleached with a 1.5% H_2_O_2_/1% KOH solution to remove pigmentation, and washed with 1x PBST. Larvae were then permeabilized with Trypsin and EDTA, washed with 1x PBST and blocked for one hour (PBS with 2% normal goat serum and 1% bovine serum albumin), and incubated with primary antibody solution (phosphorylated ERK antibody, Cell Signaling Technology, cat# 4370 and total ERK antibody, Cell Signaling Technology, cat# 4696 in 1:500 dilution) overnight. Samples were then incubated in conjugated secondary antibodies (Goat anti-rabbit IgG 594 cat# A-21125 and Goat anti-mouse IgG 488 cat# A-21121, 1:500 dilution) in 1X PBT with 1% BSA and 1% DMSO overnight. Samples were then washed with 1x PBT and spiked with 0.02% Sodium azide to prevent bacterial growth.

Larvae were embedded in 1.5% low melting point agarose in 0.68mm diameter glass capillary tubes. All images were captured using Zeiss Z.1 lightsheet confocal microscope using a 10x objective and 488 nm and 594 nm lasers. Z-stacks were obtained at 1024 x 1920 pixels with a 0.9x zoom and 2 µM step size. Larvae were imaged blind to genotype, with genotyping performed post hoc. Lightsheet stacks were converted to Dual Side Fusion .czi files using Zeiss imaging software. Dual fusion .czi files were converted to .tif, and preprocessed using a Fiji macro to generate .nrrd files with split channels. Nrrd files were then registered to a zebrafish reference brain Ref20131120pt14pl2.nrrd (zebrafishexplorer.zib.de/download) using CMTK^26,27^. Registered “warp”.nrrd images were then used to generate voxel-wise pERK activity and morphometry maps (‘map-maps’) and compared across genotypes using a MATLAB script. Region specific activity and morphometry data was then extracted using a Python script and displayed in heatmaps^28,29^. Fiji, Python, MATLAB, and R scripts can be found at github.com/melody-create/Image-Analysis and github.com/sthyme/ZFSchizophrenia/tree/master/cluster_pErk_imageprocessing.

### Calcium Imaging

*cyfip2^+/-^;mitfa^+/-^;Tg(elavl3:h2b-GCaMP6s)* adult fish were incrossed, and larval offspring were sorted at 3 dpf for *Tg(elavl3:h2b-GCaMP6s)* expression and absence of pigment (*mitfa^-/-^*). All imaging experiments took place at 5 dpf. 12 hours prior to imaging larvae were placed in a dark incubator at 29°C. Larvae were mounted dorsal side down in 2% low-melting point agarose in High Calcium Ringer’s Solution (116nM NaCl, 2.9mM KCL, 10.0 mM CaCl_2_, 5.0mM HEPES) in glass bottom petri dishes. All images were captured using a custom-built (3i) inverted spinning disk confocal microscope. The microscope contains: Zeiss Axio Observer Z1 Advanced Marina Microscope, X-cite 120LED White Light LED System, filter cube for GFP a motorized X,Y, stage, piezo Z stage, 20X Air (0.5 NA), objective, CSU-XI Spinning Disk Confocal Head with 1X camera adapter, and an Hamamatsu EMCCD camera (C9100-23B), laser stack with 405nm, 488nm, and 561nm. Fish were positioned using brightfield illumination using the eye and otolith organs as a guide, and the focal plane was set to image the largest fraction of tectal neurons positioning the tectum at the anterior image, and lateral margin of the brain at the lateral edge using the *elavl3-GCaMP5G* (slide 90/137) as reported on the Z Brain Atlas as a guide^24^. Larvae were given a 3 min rest, then the 488 nm laser was activated and images were captured at 20 fps for 30 s with consistent gain, exposure, and laser power. Following a 3 min interstimulus interval, imaging was repeated for a total of three trials per fish. All imaging and image analysis was done blind to genotype, with genotyping performed post hoc.

Images were analyzed using FIJI with the MCA plugin as previously described^30^. Cell ROIs were generated using a pretrained Cellpose model (cyto3) in combination with manually drawn ROIs based on sum projections of image stacks. A binary threshold was applied, and images were segmented using a watershed segmentation algorithm^31^. Background was subtracted using a 50-pixel rolling ball radius on all images, and baseline activity was considered to be last 5 frames of captured images. Cells in the left hemisphere were grouped by anatomical landmarks into 5 regions (pretectum, periventricular layer, neuropil, cerebellum, and hindbrain). Cells in the right hemisphere were excluded due to incomplete imaging. All data was smoothed with a gaussian filter. Data was separated into initial stimulus response (0-5s) and spontaneous activity (5-30s) and analyzed using Graphpad Prism. A threshold of 1.5 times, and 2 times the Z-score was set for a firing event in the initial stimulus response and spontaneous activity groups respectively.

### Prey Capture

All larvae were tested at 5 dpf unless otherwise stated. Larvae were randomly placed in individual wells of a 24 well plate containing 300 µL of E3 media and 20 rotifers. Plates were placed at 29°C for a total duration of five hours. After five hours, remaining rotifers were manually counted by two independent observers, and the percent rotifers consumed was calculated.

### Dark/Light Flash Assays

Prior to testing, larvae were acclimated to testing arena lighting and temperature for 30 min. Once placed on the 36 well stage in E3 media, larvae were then acclimated to dim white LED light (0.4V) for 10 min prior to dark flash onset. Larvae were then given a series of 10 ‘dark flash’ stimuli (0V) with a duration of 1s and a 30s inter stimulus interval (ISI), followed by a series of 10 ‘light flash’ stimuli (10V) with the same duration and ISI. Recordings were captured at 1000 fps and 640 x 640 px resolution using a Photron mini UX-50 camera, InfraRed illuminator, and IR diffuser. Recordings were analyzed using FLOTE as in^22,32–34^.

### Optokinetic Response

Larvae were embedded in 4% methylcellulose (28°C) oriented dorsal side up in a 35mm petri dish at 5dpf. Visual stimulus consisting of a black and white sinusoidal grating at either 15 or 30 °/s was delivered by a Samsung Galaxy Z Flip positioned 6 cm away from the petri dish and opened at a 30° angle, covering approximately 120° of the visual field as previously described^35^. Responses were recorded with a DS-Qi2 monochrome CMOS camera mounted on a Nikon SMZ25 stereo microscope at 5 frames per second. Eye movements (Δocular angle, ocular range, reflex gain) were analyzed using FIJI by thresholding to isolate the eyes and head, fitting an ellipse to each eye, and then subtracting the Δeye angle from Δhead angle to correct for any head movements. Inter eye movement responses (interocular gain, interocular concordance, interocular difference) were analyzed as previously described using MATLAB software *OKRtrack* and *OKRanalyze*^36^.

### Statistical Analysis

Statistical analyses and data visualization were performed using GraphPad Prism 10, with specific tests indicated in the results section and figure legends. MATLAB (Mathworks) scripts were used for OKR and pERK immunostaining analyses. OKR scripts are publicly available^36^, and all other MATLAB, R, and Python scripts are available at github.com/melody-create/Image-Analysis and github.com/sthyme/ZFSchizophrenia/tree/master/cluster_pErk_imageprocessing.

## Results

### *cyfip2* is required for prey capture and visually evoked startle behaviors

To determine if Cyfip2 regulates visually mediated behaviors, we performed an incross of *cyfip2^+/-^* heterozygous adults to generate *cyfip2* mutant (*cyfip2^-/-^*) and sibling (*cyfip2^+/^*) larvae. At 5 days post fertilization (dpf) we examined whether *cyfip2* mutants have a deficit in prey capture as compared to their siblings by individually incubating larvae in 24-well plates and adding 20 rotifers to each well. Larvae were left to freely swim and consume prey *ad libitum* for 5 hours, after which the number of remaining rotifers was counted. *cyfip2* mutants consumed only 4% of rotifers, whereas their siblings consumed 85% (Figure 1A, p<0.0001). Furthermore, 93% of siblings captured at least one rotifer, compared to only 25% of *cyfip2* mutants. Successful prey capture involves a complex series of behavioral bouts, requiring visual acuity and fine motor control to detect rotifers, orient body position, approach and finally strike and capture^37–39^. Such a stark deficit in the hunting ability of *cyfip2* mutants indicates a critical role for Cyfip2 in regulating visual function and/or sensorimotor integration.

**Figure 1.**
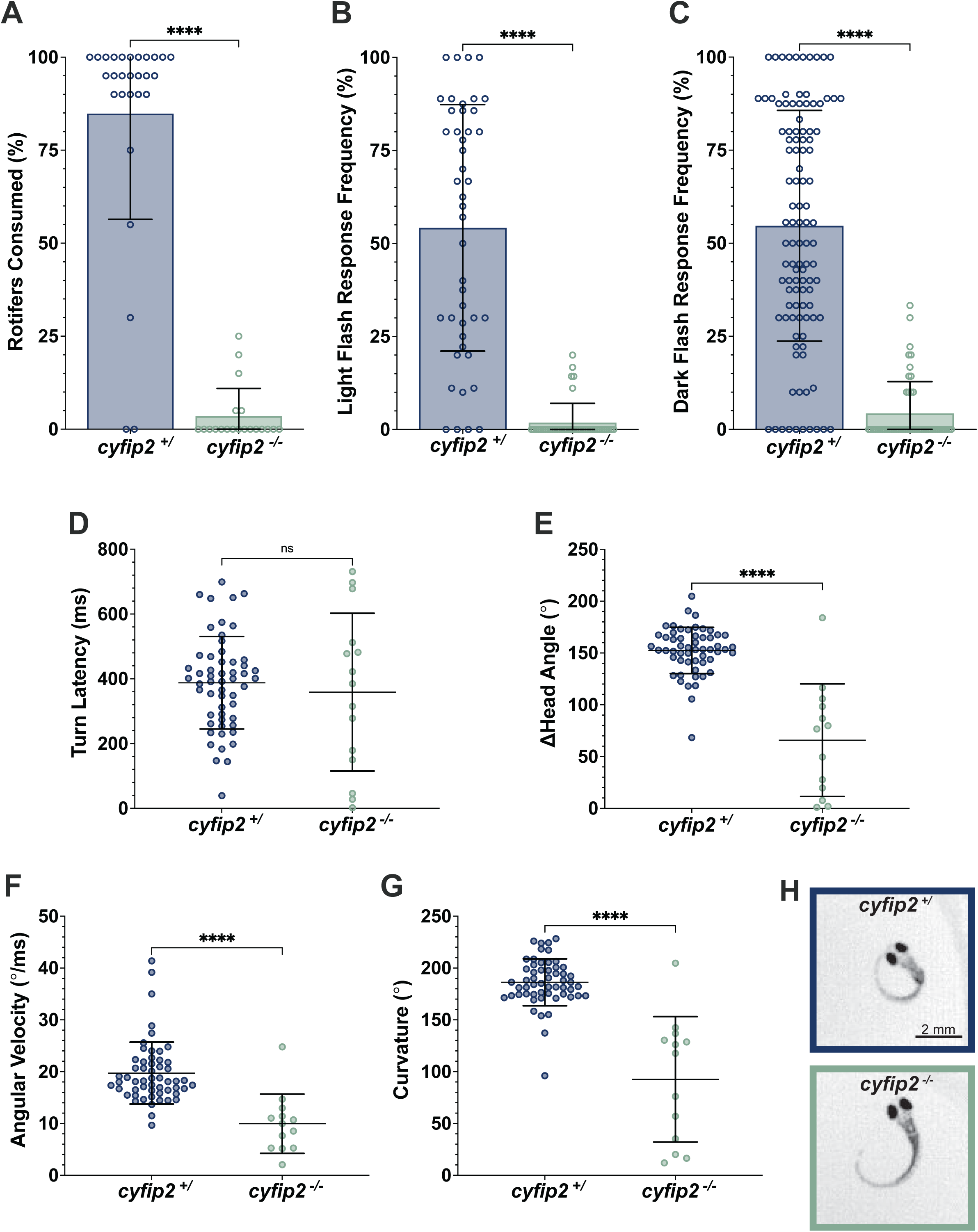
***cyfip2* mutants display stark deficits in visual behavior responses**. **A)** Percentage of rotifers consumed after 5 hours by 5 dpf siblings (*cyfip2^+/^*, blue bars, n=28) and mutants (*cyfip2^-/-^*, green bars, n=20) larvae. **B)** Response frequency to 10 light flash stimuli at 5 dpf of *cyfip2^+/^*(n=44) and *cyfip2^-/-^* (n=41). **C)** Response frequency to 10 dark flash stimuli at 5 dpf of *cyfip2^+/^*(n=107) and *cyfip2^-/-^* (n=49) larvae. **D-G)** Kinematic analysis of dark flash response latency, change in head angle, angular velocity, and curvature of *cyfip2^+/^* (n=55) and *cyfip2^-/-^* (n= 15) larvae. **H)** Representative images of the peak of an O-bend response in *cyfip2^+/^* and *cyfip2^-/-^*5 dpf larvae following dark flash stimulus. Data are presented as mean ± SD. Comparisons made using unpaired t-test, ****p < 0.0001.

To probe visual system function without requiring such a complex series of movements and feedback, we assessed dark flash and light-flash evoked startle behaviors. These sudden changes in whole-field illumination typically evoke large angle turns called O-bends^33,34,40^. 5 dpf larvae were placed in individual wells of a custom, IR-illuminated 36-well acrylic grid placed beneath a high-speed camera with an IR filter^23,41,42^. LED illumination of the grid was set to low (0.4V) while the larvae habituated for 10 minutes. Then larvae were then given a series of ten, 1 sec dark flashes (0V) separated by a 30 sec inter-stimulus interval (ISI), followed by ten, 1 sec light flashes (10V) separated by a 30 sec ISI. Responses were recorded at 1,000 frames per second, and larval behavior was tracked and analyzed in an unbiased, high-throughput manner using FLOTE^32,34^. We observed that *cyfip2* mutants are almost completely unresponsive to both light flash and dark flash stimuli (Figures 1B and 1C). On average, *cyfip2* mutants responded to only 4% of dark flash stimuli, as compared to a 55% response frequency of siblings (p<0.0001), a 92% decrease in response frequency of *cyfip2* mutants. Additionally, 76% of *cyfip2* mutants never responded to dark flash stimuli, compared to just 9% of siblings. Responses to light flash stimuli showed a similar pattern. On average, *cyfip2* mutants responded to 2% of light flash stimuli, compared to 54% response frequency of siblings (p<0.0001), a 96% reduction in response frequency of *cyfip2* mutants. Furthermore, 88% of *cyfip2* mutants were never responded to light flash stimuli, compared to 9% of their siblings.

Because *cyfip2* mutants display altered kinematics of acoustically-evoked startle responses^22,23^, we next wanted to determine if the subset of *cyfip2* mutants (24%) that did respond to dark flash stimuli had normal O-bend kinematics as compared to their siblings. We compared the average values of response latency, change in head angle, angular velocity, and curvature (summation of head and tail angle) of siblings and responsive *cyfip2* mutants. We found that while the turn latency of *cyfip2* mutants is not significantly different than their siblings (Figure 1D, p=0.5577), *cyfip2* mutants performed shallower turns with reduced changes in head angle, angular velocity, and curvature (66°, 10°/ms, and 93°, respectively) as compared to their siblings (152°, 20°/ms, and 186°; Figures 1E-G, p<0.0001). This indicates that of the small proportion of *cyfip2* mutants that do respond to dark flash stimuli, these responses display a weaker activation of the trunk musculature, resulting in slow and shallow turning movements (Figure 1H).

### *cyfip2* is dispensable for the Optokinetic Response

The severe deficits in prey capture and responses to whole field illumination changes in *cyfip2* mutants led us to ask whether defects in retinal phototransduction could account for all of these phenotypes, particularly as defects in zebrafish retinal lamination^11^ and mouse retinal ganglion cell (RGC) function^43^ have been observed in *cyfip2* knockout animals. To determine if *cyfip2* mutants have intact retinal phototransduction, we tested the optokinetic response (OKR), an innate highly conserved, innate behavior that stabilizes images on the retina to enable high resolution vision^44,45^. Unlike prey capture and O-bend responses, this behavior does not require coordinated activation of hindbrain and spinal locomotor circuits, and it is independent of the optic tectum^46^, the primary recipient of RGC inputs, so OKR defects in *cyfip2* mutants would implicate the retina as the likely source of all visual deficits. To examine differences in the OKR of *cyfip2* mutants and siblings, we embedded 5 dpf larvae in methylcellulose and provided a sinusoidal grating stimulus encompassing 120° of the visual field as previously described^35^ while recording eye movements from above. The OKR assay consisted of three phases; alternating, unidirectional (right), and unidirectional (left) performed at two grating speeds, ‘slow grating’ (15°/s) followed by ‘fast grating’ (30°/s) (Figure 2A). When we examined the change in ocular angle of the left and right eyes throughout the experiment, we found that *cyfip2* mutants are nearly as capable of tracking both fast and slow grating stimuli as their siblings (Figures 2A,B). We did find that *cyfip2* mutants have a deficit when tracking leftward motion with the left eye (Figure 2C), and rightward motion with the right eye (Figure 2D), suggesting a defect in the function of either the lateral rectus muscle or the abducens nerve (cranial nerve VI) that innervates it, allowing for outward eye movement (Supplementary Video S3,4)^47,48^.

**Figure 2.**
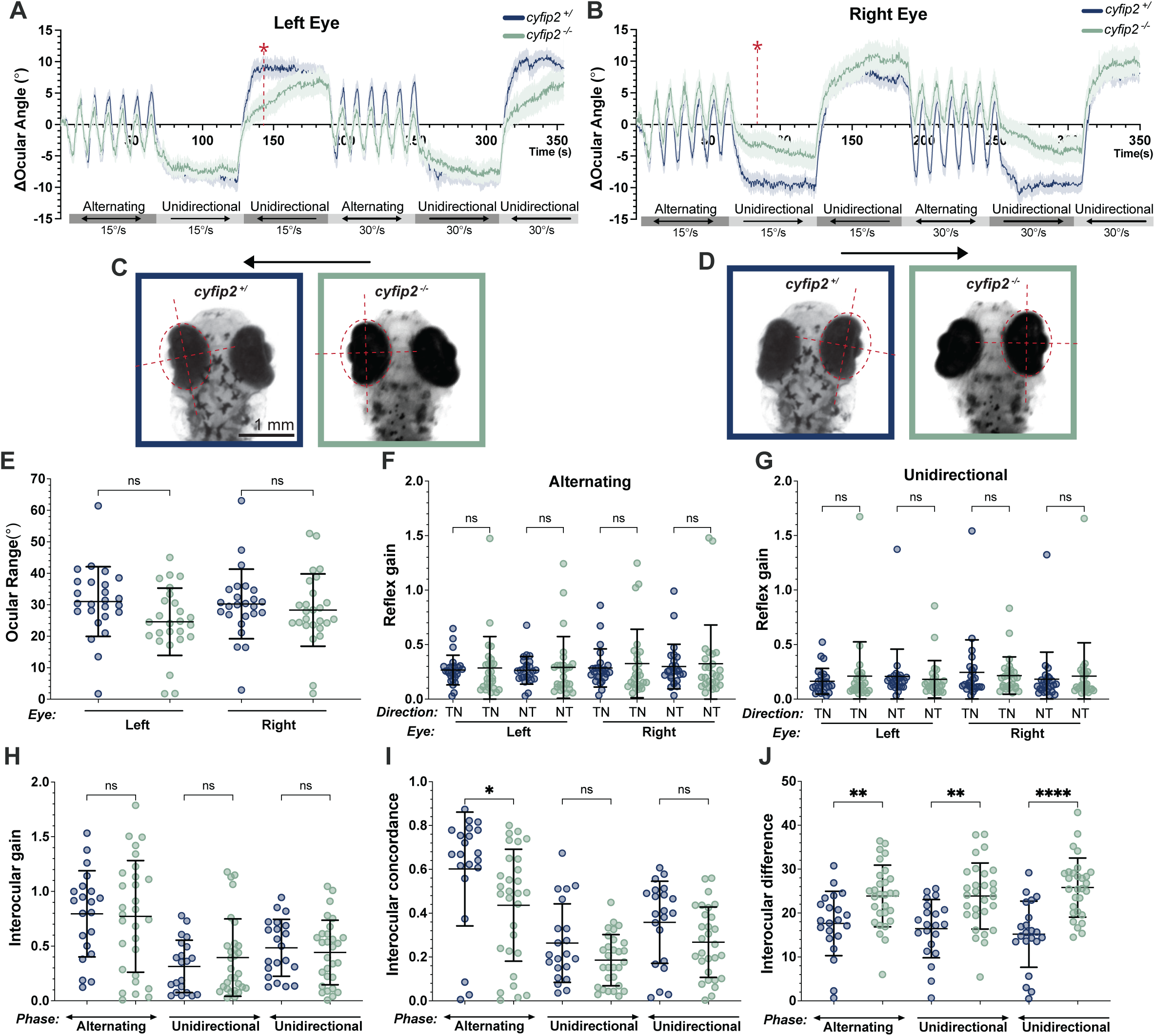
*cyfip2* mutants have subtle defects in the Optokinetic Response. Quantification of optokinetic response to 99% contrast sinusoidal grating moving in either alternating or unidirectional pattern at a velocity of 15°/s or 30°/s. **A)** Mean trace of ocular angle of left eye over time of 5 dpf *cyfip2^+/^* (blue line, n=25) and *cyfip2^-/-^* (green line, n=26) larvae (mean ± SEM). Red star indicates the timepoint of images shown in **C**. **B)** Mean trace of ocular angle of right eye over time of *cyfip2^+/^* (n=25) and *cyfip2^-/-^* (n=26) larvae (mean ± SEM). Red star indicates the timepoint of images shown in **D. C,D)** Representative images showing defect in outward eye movements of *cyfip2^-/-^* during the left (C) and right (D) unidirectional 15°/s stimulus. **E)** Total ocular range of left and right eye of *cyfip2^+/^*(n=25) and *cyfip2^-/-^* (n=26) larvae (mean ± SD). Reflex gain of left or right eye in nasotemporal (NT) or temporonasal (TN) direction during alternating **F)** and unidirectional **G)** phase of 15°/s sinusoidal grating of *cyfip2^+/^*(n=25) and *cyfip2^-/-^* (n=26) larvae (mean ± SD). Between-eye measurements of interocular gain **H)** interocular concordance **I)** and interocular difference **J)** during alternating and unidirectional phase of sinusoidal grating at velocity of 15°/s and 30°/s of *cyfip2^+/^* (n=21) and *cyfip2^-/-^* (n=29) larvae (mean ± SD). Comparisons made using 1-way ANOVA with Šídák’s multiple comparisons test, * p < 0.05, **p < 0.01, ****p < 0.0001.

Despite this defect in outward motion of each eye, we found no difference in the total ocular range of either left or right eye in *cyfip2* mutants and their siblings (Figure 2E). Additionally, we found no differences in the reflex gain (ratio of eye angular velocity to stimulus angular velocity) between *cyfip2* mutants and siblings in the ‘slow grating’ alternating or unidirectional phases for either left or right eye, regardless of stimulus direction (Figure 2F,G). This held true for the ‘fast grating’ as well (Supplementary Fig. S1). This indicates that *cyfip2* mutants are capable detecting the visual grating stimulus and responding with appropriate eye tracking and image stabilization movements. Finally, we examined the relationship between the movement of both eyes and found no difference in interocular gain (ratio of angular displacement of right eye to left eye) between *cyfip2* mutants and siblings, indicating that the eyes are moving at the same speed (Figure 2H). We also found no difference in interocular concordance (proportion of eye movements with the same direction of displacement) of *cyfip2* mutants and siblings, indicating that the eyes are moving together (Figure 2I). We did observe a significant increase in the interocular difference (difference between eye angles) of *cyfip2* mutants as compared to siblings (Figure 2J, p=0.0079, p=0.0011, p<0.0001), reflecting the deficit in outward movement of each eye noted above.

Overall, our data show that *cyfip2* mutants have relatively subtle defects in the OKR, limited to the extent of outward eye movements. That mutants are able to detect and track the movement of the grating stimulus indicates that retinal phototransduction remains intact. Thus, *cyfip2* is likely dispensable for retinal function, and the loss of prey capture and dark flash and light-flash responses suggest that *cyfip2* has more specific roles in the brain to enable visual sensorimotor integration.

### Loss of *cyfip2* reduces optic tectum volume and basal forebrain and midbrain activity

To identify potential sites in the brain where *cyfip2* acts to regulate visually-driven behavioral responses, we measured brain-wide activity and morphometry using a well-established Mitogen Activated Protein “MAP-mapping” immunostaining approach that compares the ratio of phosphorylated ERK (pERK) to total ERK (tERK) as a readout of neural activity^24–26^. 5 dpf larvae were allowed to swim freely in 10 cm petri dishes for 30 min prior to fixation to examine the basal activity state of the brain. Following staining, fish were imaged by lightsheet microscopy with genotyping performed post-hoc. Compared to wild-type and heterozygous siblings, *cyfip2* mutants have decreased activity within the forebrain (olfactory bulb, subpallium, pallium) and midbrain (habenula, dorsal thalamus, ventral thalamus, pretectum, and tectum neuropil) (Figure 3A, C). And by analyzing brain morphometry using tERK labeling we found that this broad decrease in activity within the midbrain correlates with a pronounced decrease in volume of the tectum neuropil (Figure 3B, D). A complete list of all 294 brain regions analyzed for morphometry and activity are included in (Supplementary Figure S2, S3) decrease in both neural activity and volume within the optic tectum could account for *cyfip2* mutants’ deficits in visual behavior responses, particularly as the intact OKR is independent of the tectum^46,49,50^. To our surprise, based on previous findings of hyperactivity within the acoustic startle circuit of *cyfip2* mutants^22,23^, we did not observe any differences in basal neural activity or morphology in the hindbrain, suggesting that mutants’ acoustic hypersensitivity does not emerge from altered basal activity but rather from heightened stimulated activity. Together, these results indicate that the optic tectum is the likely site where *cyfip2* acts to mediate visual behaviors.

**Figure 3.**
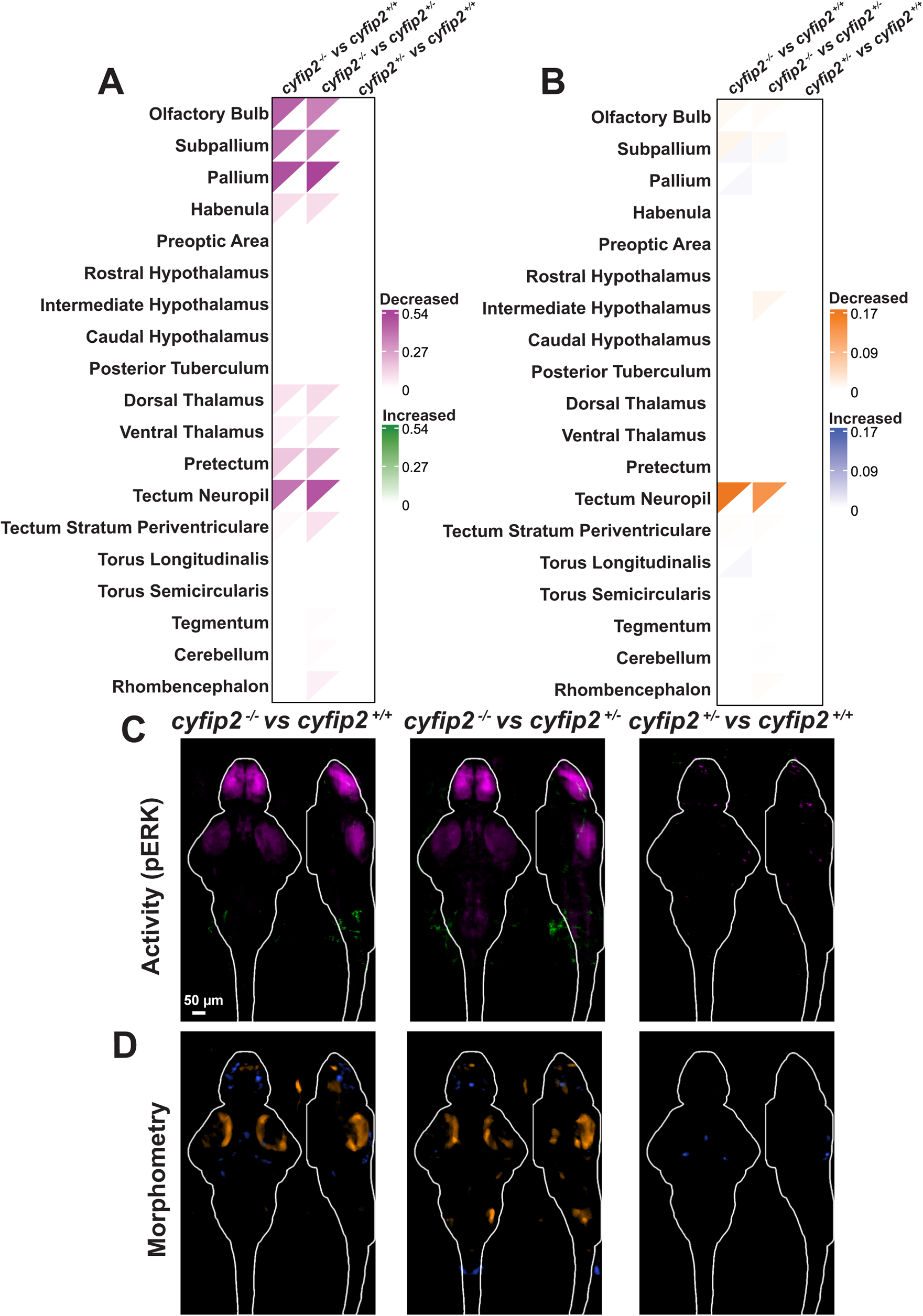
*cyfip2* mutants display both decreased activity and volume in the optic tectum. Heatmap of changes in **A)** activity and **B)** morphometry across 19 brain regions in 5 dpf larvae, comparing all three *cyfip2* genotypes to one another. Decreases in activity (magenta) and morphometry (orange) are shown in the upper left corner and increases in activity (green) and morphometry (blue) in the lower right corner. Dorsal (Z plane, left) and lateral (X-Y plane, right) sum projections of significant differences in **C)** activity and **D)** morphometry between genotypes, with the whole brain outlined in white.

### Loss of *cyfip2* dampens visually evoked neural activity without altering firing probability or overall response patterns across brain regions

To more directly link the substantial decrease in unstimulated basal neural activity in *cyfip2* mutants (Figure 3) to their deficits in visual behavior responses, we sought to determine if visual stimulus-evoked activity was also disrupted by loss of *cyfip2*. We used the pan-neuronal, nuclear-localized GCaMP transgenic line *Tg(elavl3:h2b-GCaMP6s)* in the pigment mutant *mitfa-/-* background to compare neural activity of *cyfip2* wildtype (n=4), *cyfip2* heterozygous (n=4), and *cyfip2* mutant (n=4) larvae following a light stimulus. We recorded 30 sec of GCaMP6s fluorescence from a single focal plane containing the tectal circuits of interest^24^ with a spinning disk confocal microscope, using the light from the 488nm imaging laser as a visual stimulus. Following a 3-minute interstimulus interval, this assay was repeated for a total of 3 trials per fish. Images were registered to correct for motion, and cells were segmented using the FIJI CellPose plugin^30,51^ with manual supervision. Identified cells were grouped by anatomical landmarks into 5 regions (pretectum, periventricular layer, neuropil, cerebellum, and hindbrain) with an average of 136 total cells analyzed per fish (Supplementary Figure S6, A). We calculated Z-scores for the change in GCaMP6s fluorescence for each cell across the three trials. We considered the first 5 seconds of activity to be the immediate response to the light stimulus (Figure 4) and the following 25 seconds to reflect more spontaneous activity and habituation to the stimulus (Supplementary Figures S5, S6).

**Figure 4:**
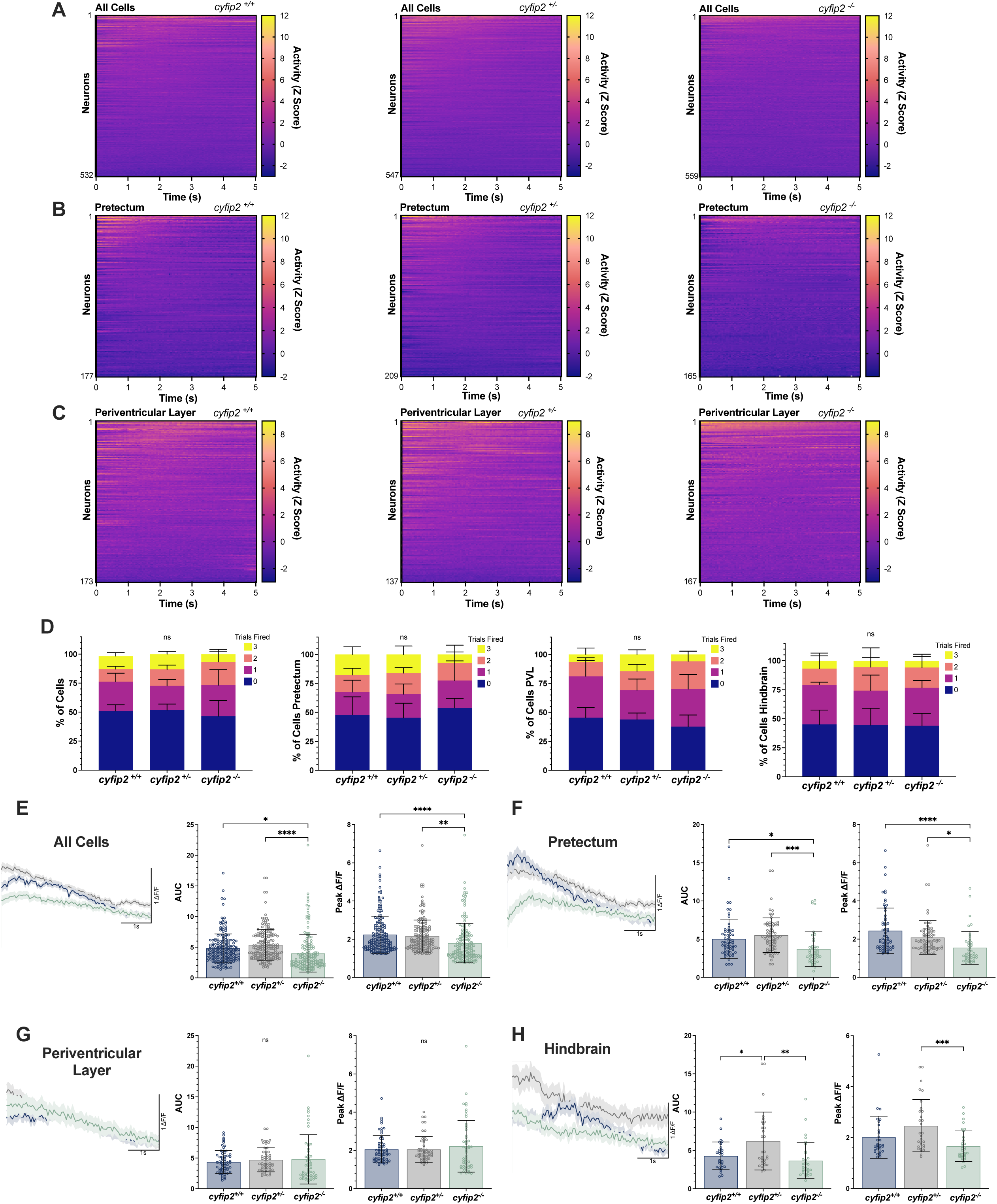
**Loss of Cyfip2 causes dampened neuronal activation following light stimulus**. Normalized heatmaps depicting neural activity in **A)** all neurons, **B)** Pretectum neurons, and **C)** Periventricular Layer neurons during the first 5 seconds following a blue light (488nm) stimulus in *cyfip2^+/+^*, *cyfip2^+/-^* and *cyfip2^-/-^*. **D)** The percentage of neurons that fire (Z score >1.5) over the course of three successive trials, 3 times (yellow), 2 times (orange), 1 time (magenta), and 0 times (dark purple), in all cells, the Pretectum, the Periventricular Layer (PVL) and the Hindbrain across all three genotypes. Chi square test was used to determine significance. **E-H)** The average ΔF/F traces for all firing events, the area under the curve (AUC) of the ΔF/F trace or each active neuron, and the peak ΔF/F value for *cyfip2^+/+^* (blue), *cyfip2^+/-^* (grey) and *cyfip2^-/-^* in **E)** all neurons, **F)** the Pretectum, **G)** the Periventricular Layer, and **G)** the Hindbrain. Data are presented as mean ± SEM for the average ΔF/F traces, and mean ± SD for all other data. Comparisons made using 1-way ANOVA with Tukey’s multiple comparisons test, * p < 0.05, **p < 0.01, ***p < 0.001, ****p < 0.0001.

Following the light stimulus, we observed a wide range of activity patterns across neurons within each genotype, with many cells responding immediately, some with later onset, and many not responding at all (Figure 4A-C). To determine if *cyfip2* regulates these patterns of activity, we performed a Principal Component Analysis (PCA) and found no clear segregation of genotypes (Supplementary Figure S4), indicating that *cyfip2* does not play a role in the overall pattern of neuronal responses. We then asked whether *cyfip2* regulates the likelihood of neuronal firing, and so we calculated the percentage of cells that fired during 3, 2, 1, or 0 trials across: 1) all cells, 2) the pretectum, 3) the periventricular layer, and 4) the hindbrain, with a firing event defined as a Z-score greater than 1.5 (Figure 4D). There were no significant differences in the percentage of cells that fired between wild-type, *cyfip2* heterozygous, or *cyfip2* mutant fish in any brain region, and across all cells analyzed. However, while neurons in *cyfip2* mutants were no less likely to fire, the change in fluorescence of *cyfip2* mutant neurons was dampened compared to wild-type and heterozygous neurons (Figure 4E). Across all cells, during trials in which they fired (Z-score > 1.5) we observed significant decreases in the area under the curve (AUC) of each ΔF/F trace (p=0.0109) and in the peak ΔF/F signal in *cyfip2* mutants compared to wild-type and heterozygous fish (p<0.0001). This decrease in all cells’ activity was largely explained by the decrease in fluorescence signal we observed in the pretectum, with a decrease in AUC (p=0.0120) and peak ΔF/F signal (p<0.0001) (Figure 4F). There were no significant differences in ΔF/F in the periventricular layer across genotypes (Figure 4G). We did observe an interesting increase in hindbrain activity in *cyfip2* heterozygotes (Figure 4H), but it is unclear how meaningful this difference is since no behavioral deficits in *cyfip2* heterozygotes have been reported^22,23^. Overall, while *cyfip2* mutant neurons are no more likely to fire following a light stimulus, when they do fire their signal is dampened, particularly for neurons in the pretectum and tectum neuropil (Supplementary Figure S6).

Finally, we wanted to determine if *cyfip2* mediates spontaneous activity following an evoked stimulus, which we considered to be the period of 5 to 30 seconds following the light stimulus (Supplementary Figure S7). After performing a PCA, we found no clear segregation of genotypes (Supplementary Figure S4), confirming that *cyfip2* does not regulate activity patterns across neurons. However, in contrast to the decreases in activity observed in *cyfip2* mutants during the immediate response to the light stimulus (Figure 4), we observed significant increases in the AUC of both the ΔF/F trace (p<0.001) and in the peak ΔF/F signal in *cyfip2* mutants compared to wild-type and heterozygous fish (p<0.001) across all cells during trials in which they fired (Z-Score > 2.5) (Supplementary Figure S8). This increase is driven by a delayed surge in activity of cells in the periventricular layer in *cyfip2* mutants, arising 20 seconds after the light stimulus (Supplementary Figure S8, D) and leading to a significant increase in the AUC of the ΔF/F trace (p<0.0001) but not in the peak ΔF/F signal in *cyfip2* mutants compared to wild-type and heterozygous fish (p=0.3138). This delayed onset of activity within the intrinsic neurons of the tectum is reflective of the dysregulated excitability observed more broadly in *cyfip2* mutants^22,23,52–56^.

### *cyfip2* mediates early developmental assembly of visual behavior circuits through Rac1-dependent actin remodeling, independent of FMRP-dependent translational regulation

To determine the relevant developmental windows and molecular mechanisms that enable Cyfip2 to regulate visual behavior, we temporally controlled Cyfip2 expression using the transgenic line *Tg(hsp70:cyfip2-GFP)*^22^. Following a 40 min heat shock, expression of Cyfip2-GFP peaks between 2-12 hours post heatshock before tapering to undetectable levels by 18 hours post heatshock^57^, allowing for a 10-hour window of abundant Cyfip2 expression in the CNS. Larvae were heatshocked at 38°C for 40 minutes at either 0, 1, 2, 3, 4 or 5 dpf or with a multi-day heatshock protocol on 1,2,3 and 4 dpf, and then allowed to recover until dark flash/light flash response testing at 5 dpf. *cyfip2* mutants that were heatshocked at 1 dpf but did not express the heatshock-inducible transgene displayed the same severe deficits in dark flash and light flash responses compared to their siblings (Figure 5A-B), and transgene-expressing but non-heatshocked larvae showed the same pattern (Supplementary Figure S9,1E-F), indicating no effect of either the heatshock or the transgene alone. Heatshock induction of Cyfip2-GFP expression at 1 dpf, 2 dpf, or 1-4 dpf all significantly increased *cyfip2* mutants’ dark flash response frequencies, with the 1 dpf heatshock providing the strongest rescue (43%) compared to non-transgenic *cyfip2* mutant controls (4%) (Figure 5A, p<0.0001). Furthermore, while 88% of non-transgenic *cyfip2* mutants were unresponsive to dark flash stimuli, inducing expression of Cyfip2 at 1 dpf completely eliminated non-responding *cyfip2* mutants. Inducing expression of Cyfip2 at 2 dpf reduced the nonresponders to 11%, while the multi-day heatshock in which Cyfip2 was repeatedly induced on days 1-4 reduced nonresponders to 35% (Supplementary Figure S9). We observed a similar pattern for rescue of light flash response frequency, with heatshock at 1 dpf providing the strongest rescue (46% mean response frequency, 0% nonresponsive) (Figure 5B, p<0.0001), followed by 1-4 dpf (25% mean, 24% nonresponsive) (Figure 5B, p=0.0004) and 2 dpf (22% mean, 22% nonresponsive) (Figure 4B, p=0.0216). Inducing expression of Cyfip2 at 0, 3, 4, or 5 dpf did not significantly affect either dark flash or light flash responsiveness of *cyfip2* mutants. Together, these data demonstrate that Cyfip2 does not act acutely but rather functions during early neural development (30 to 50 hpf) to regulate visual responses to changes in whole-field illumination.

**Figure 5.**
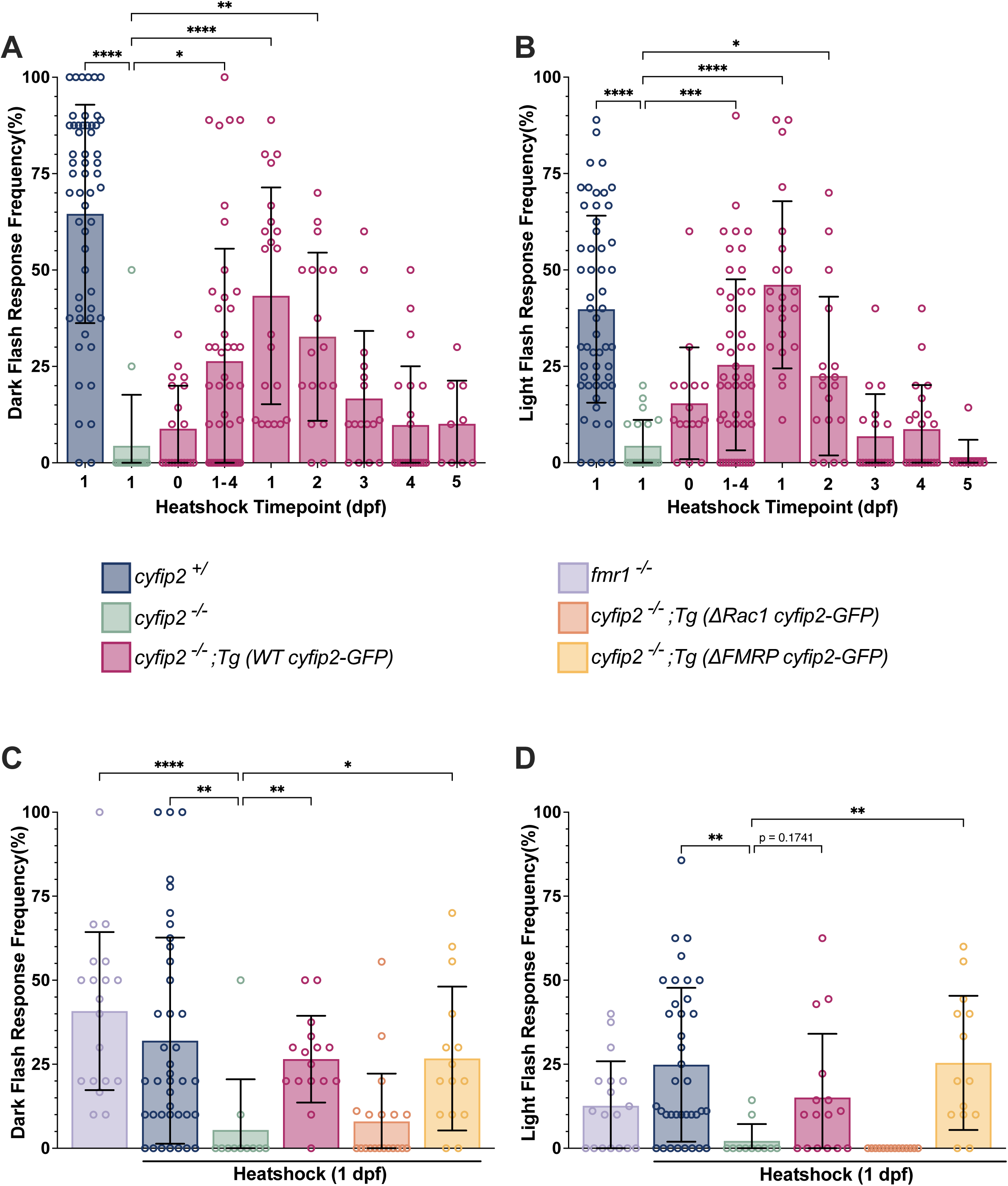
*cyfip2* developmentally regulates visual behavior through Rac1-dependent actin remodeling. **A)** Response frequency of 5 dpf *cyfip2^+/^* (blue bars, n=55), *cyfip2^-/-^*(green bars, n=17), and *cyfip2^-/-^;Tg(WT cyfip2-GFP)* (pink bars) larvae to 10 dark flash stimuli following heatshock at 12 hpf (0 dpf; n=20), 30 hpf (1 dpf; n=21), 54 hpf (2 dpf; n=18), 78 hpf (3 dpf; n=16), 96 hpf (4 dpf; n=21), 120 hpf (5 dpf; n=9), and 30,54,78, and 102 hpf (1-4 dpf; n=46). **B)** Response frequency to 10 light flash stimuli at 5 dpf after heatshock of *cyfip2^+/^*(n=54), *cyfip2^-/-^*(n=21), and *cyfip2^-/-^;Tg(WT cyfip2-GFP)*, at 0 dpf (n=16), 1 dpf (n=22), 2 dpf (n=18), 3 dpf (n=18), 4 dpf (n=21), 5 dpf (n=10), and 1-4 dpf (n=50). **C)** Response frequency to 10 dark flash stimuli at 5 dpf of *fmr1^-/-^* (purple bar, n=19), after heatshock at 30 hpf of *cyfip2^+/^* (n=39), *cyfip2^-/-^* n=11), *cyfip2^-/-^;Tg(WT cyfip2-GFP)* (n=16), after heatshock at 30 hpf of *cyfip2^-/-^;Tg(*Δ*Rac1 cyfip2-GFP)* (orange bar, n=19), and *cyfip2^-/-^;Tg(*Δ*FMRP cyfip2-GFP)* (yellow bar, n=15). **D)** Response frequency to 10 dark flash stimuli at 5 dpf of *fmr1^-/-^* (n=18), after heatshock at 30 hpf of *cyfip2^+/^* (n=37), *cyfip2^-/-^* (n=11), *cyfip2^-/-^;Tg(WT cyfip2-GFP)* (n=16), after heatshock at 30 hpf of *cyfip2^-/-^;Tg(*Δ*Rac1 cyfip2-GFP)* (n=14), and *cyfip2^-/-^;Tg(*Δ*FMRP cyfip2-GFP)* (n=14). Data are presented as mean ± SD. Comparisons made using Kruskal-Wallis test with Dunn’s multiple comparisons correction. * p < 0.05, **p < 0.01, ***p < 0.001, ****p < 0.0001.

To determine if Cyfip2 regulates the development of visual behavior circuitry through actin remodeling in its role as a member of the Wave Regulatory Complex (WRC) or by regulating mRNA translation through interactions with FMRP and eIF4E, we used heatshock inducible transgenic lines with point mutations in Cyfip2’s Rac1 *Tg(hsp70:cyfip2-(C179R(*Δ*Rac1))-EGFP*) and FMRP *Tg(hsp70:cyfip2-(K723E(*Δ*FMRP))-EGFP)* binding domains^23^, hereafter referred to as *Tg(*Δ*Rac1 cyfip2-GFP)* and *Tg(*Δ*FMRP cyfip2-GFP)*. Previous studies have established that C179R and K723E amino acid substitutions prevent Cyfip2 from binding with Rac1 and FMRP respectively, and therefore inhibit its ability to interact in either of these pathways^12,14,20^. As both *Tg(*Δ*Rac1 cyfip2-GFP)* and *Tg(*Δ*FMRP cyfip2-GFP)* produced weaker GFP expression following a 40 minute heatshock at 38°C compared to *Tg (WT cyfip2-GFP)*^23^ (Supplementary Figure S9, A-C) we reduced the heatshock duration of *Tg (WT cyfip2-GFP)* to 25 minutes to provide comparable levels of Cyfip2-GFP, ΔRac1 Cyfip2-GFP, and ΔFMRP Cyfip2-GFP (Supplementary Figure S9). This 25 min heatshock of *Tg (WT cyfip2-GFP)* at 1 dpf was sufficient to rescue dark flash responsiveness and improved light flash responsiveness (Figure 5C,D, Supplementary Figure S9I, J).

We found that inducing expression of *cyfip2^-/-^;Tg(*Δ*Rac1 cyfip2-GFP)* failed to improve either dark flash (8%) or light flash (0%) responsiveness compared to non-transgenic *cyfip2* mutant controls (5% and 2% respectively) (Figure 5C,D, p>0.9999, p<0.9999). Therefore, Cyfip2 must interact with the WRC to regulate the development of visual behavior circuitry. However, we found that inducing expression of *Tg(*Δ*FMRP cyfip2-GFP)* rescued both dark flash (27%) and light flash (25%) responsiveness (Figure 5C,D, p=0.0150, p=0.0038) and reduced the percentage of non-responders substantially (Supplementary Figure S9I,J). Therefore, Cyfip2 acts independently of FMRP/eIF4E during this developmental window to regulate visual behavior responses. To further validate this finding, we tested whether FMRP is required for responses to whole-field illumination changes by measuring dark flash and light flash response frequency in *fmr1* mutant larvae. *fmr1* mutants had similar dark flash and light flash response frequencies to *cyfip2* wildtypes and heterozygotes (Figure 5C,D), confirming that interactions between FMRP and Cyfip2 are not required for these visually-mediated behaviors.

We previously observed that while the majority of *cyfip2* mutants do not respond to dark flash stimuli, the small portion that do respond display slower, shallower responses (Figure 1B-H). And so, we sought to determine the molecular requirements for *cyfip2*’s role in regulating visual response kinematics using the same heatshock approach. We found no significant differences in turn latency for any group (Figure 6A), but heatshock induction of *Tg(WT cyfip2-GFP)* at 1 dpf increased the change in head angle, angular velocity, and curvature of *cyfip2* mutants (Figure 6 B,D, p=0.0089, p=0.0791, p=0.0062, respectively). In contrast, heatshock-induced expression of *Tg(*Δ*Rac1 cyfip2-GFP)* did not improve the change in head angle, angular velocity, or curvature (Figure 6B-E), consistent with its failure to rescue dark flash and light flash responsiveness (Figure 5 C,D). We also found that induction of *Tg(*Δ*FMRP cyfip2-GFP)* does restore changes in head angle (Figure 6B, p<0.0001), angular velocity (Figure 6C, p=0.0004), and tail angle (Figure 6D, p<0.001). These findings further confirm that Cyfip2 acts through Rac1 and the WAVE Regulatory Complex but not through FMRP/eIF4E to regulate not only the initiation but also the kinematic performance of responses to changes in whole field illumination.

**Figure 6.**
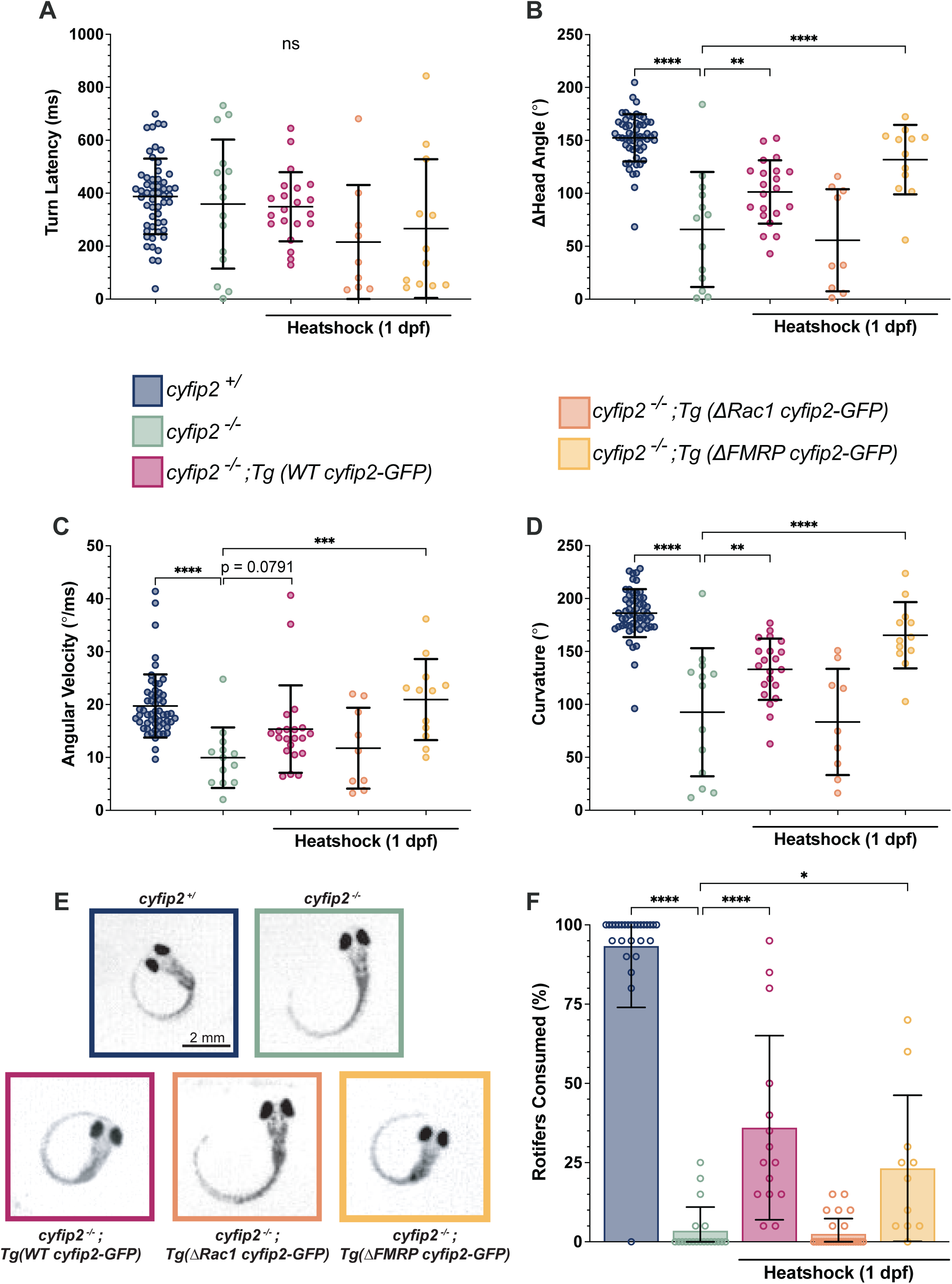
Developmental expression of Cyfip2 rescues O-bend kinematics and prey capture responses. A-D) Kinematic analysis of dark flash responses of *cyfip2^+/^* (blue dots, n=55) and *cyfip2^-/-^* (green dots, n= 15) at 5dpf, and *cyfip2^-/-^;Tg(WT cyfip2-GFP)* (pink dots, n=21), *cyfip2^-/-^;Tg(*Δ*Rac1 cyfip2-GFP)* (orange dots, n=9), and *cyfip2^-/-^;Tg(*Δ*FMRP cyfip2-GFP)* (yellow dots, n=12) at 5 dpf after heatshock at 38°C at 30 hpf (1 dpf), as analyzed by FLOTE software. **E)** Representative images of the peak of the O-bend response of *cyfip2^+/^*, *cyfip2^-/-^*, *cyfip2^-/-^;Tg(WT cyfip2-GFP*, *cyfip2^-/-^;Tg(*Δ*Rac1 cyfip2-GFP )*, and *cyfip2^-/-^;Tg(*Δ*FMRP cyfip2-GFP)* larvae following dark flash stimulus. **F)** Percentage of rotifers consumed after 5 hours by 5 dpf *cyfip2^+/^* (n=28), *cyfip2^-/-^*(n=20), *cyfip2^-/-^;Tg(WT cyfip2-GFP)* (n=15), *cyfip2^-/-^;Tg(*Δ*Rac1 cyfip2-GFP )* (n=28), and *cyfip2^-/-^;Tg(*Δ*FMRP cyfip2-GFP)* (n=11). Data are presented as mean ± SD. Comparisons made using 1-way ANOVA with Šídák’s multiple comparisons test, * p < 0.05, **p < 0.01, ***p < 0.001, ****p < 0.0001.

Finally, to determine if developmental expression of Cyfip2 is sufficient to enable more demanding visually-mediated behaviors, we heatshocked fish for 40 min at 1 dpf and assessed prey capture at 5 dpf as previously described (Figure 1). Expression of *Tg(WT cyfip2-GFP)* significantly improved rotifer capture in *cyfip2* mutants from 4% to 36% of rotifers (Figure 6F, p<0.0001). Additionally, 100% of *Tg(WT cyfip2-GFP)*-expressing *cyfip2* mutants were able to capture at least one rotifer, a stark improvement from the 25% of non-transgenic mutants. And consistent with our findings for dark flash and light flash responsiveness (Figure 5) and kinematics (Figure 6 A-E), we found that inducing expression of *Tg(*Δ*Rac1 cyfip2-GFP)* failed to rescue prey capture (2.5%) (Figure 6F, p=0.8662), whereas expression of *Tg(*Δ*FMRP cyfip2-GFP)* significantly improved prey capture (23%) (Figure 6F, p=0.0213). These results demonstrate that Cyfip2 mediates sensorimotor integration of visual input through Rac1-dependent actin remodeling during early neural development.

## Discussion

Cytoplasmic Fragile X Messenger Ribonucleoprotein (FMRP) Interacting Protein 2 (Cyfip2) has well-established and conserved roles in axon guidance, synapse formation, and circuit maturation^8,13,58–61^. In humans, variants in CYFIP2 cause severe neurodevelopmental challenges, including developmental delay, early infantile epileptic encephalopathy, intellectual disability, and visual impairment (collectively termed Developmental and Epileptic Encephalopathy, DEE65)^52–56^, highlighting the importance of understanding Cyfip2’s functions. The severity of these phenotypes varies with mutation location, with mutations at p.Arg87 and p.Asp724 hotspots producing profound deficits that include visual abnormalities (divergent strabismus, cortical vision loss, optic nerve atrophy), microcephaly, truncal hypotonia, severe intellectual disability, feeding and swallowing difficulties, hyperkinetic behavior, and disturbed sleep^55,56^. In zebrafish, mutations in *cyfip2* disrupt the dorsoventral pattern of retinotectal axon projections to the optic tectum and alter the laminar positioning of amacrine and retinal ganglion cell (RGCs) within the retina^8–11,43^. These findings indicate a critical role for Cyfip2 in visual circuit organization, but the developmental timing and cellular and molecular pathways by which Cyfip2 acts to enable visual function and behavioral responses to visual stimuli had not been examined. Here we used a combination of behavioral assays including prey capture, dark flash and light flash stimuli, and the optokinetic response (OKR), along with brain-wide functional imaging and conditional rescue experiments to determine Cyfip2’s roles in the visual system.

We show that *cyfip2* mutant zebrafish larvae exhibit severe impairments in prey capture and visual startle responses, despite preserved retinal phototransduction as evidenced by an intact OKR. Mutants also exhibit reduced basal neural activity in specific brain regions including the primary visual processing center, the optic tectum. Developmental rescue experiments revealed that *cyfip2* regulates the development of visual circuits during a critical window between 30-50 hpf via Rac1-dependent actin remodeling. These findings support and expand upon previous studies showing a role for Cyfip2 in retinal ganglion cell (RGC) axon guidance^8,11,12^ by defining the spatial, temporal, and molecular requirements for Cyfip2 to enable functional visual sensorimotor integration.

*cyfip2* mutants have pronounced deficits in their ability to capture prey (Figure 1), a visually-driven behavior requiring intact sensorimotor integration and higher order processing^62^. This deficit could arise from disruptions to one or more of a broad set of neural circuits for phototransduction, coordinated eye movement and tracking, premotor processing, and hindbrain coordination that together provide the ability to detect, interpret, and respond to fast-moving prey^37–39^. To assess the function of a more specific subset of circuits, we assessed visual startle responses to changes in whole-field illumination, observing severely reduced responses to dark and light flashes in *cyfip2* mutants. These “O-bend” startle responses depend on intact retinal function^63^, are mediated by visual sensory areas of the midbrain, diencephalon, and cerebellum^64^, and are not reliant on the Mauthner circuit that drives the acoustic startle response, but rather they require ventromedial reticulospinal neurons that drive spontaneous turning behavior^65,66^. Our results thus point to disruptions in one or more of these specific sensorimotor pathways. To further narrow down the possible circuits impacted by loss of *cyfip2*, we examined the OKR, a swim-independent assay that measures larval eye-tracking ability, a crucial component of effective prey capture^35,67^. *cyfip2* mutants displayed an intact OKR, indicating functional retinal phototransduction and implicating downstream sensorimotor integration as the major site of deficit, rather than stimulus detection.

Cyfip2 exerts its through two molecular pathways: translational regulation of target mRNAs via interactions with binding partners FMRP and eukaryotic translation initiation factor 4E (eIF4E)^20,21^, and modulation of actin dynamics as a component of the WASP-family verprolin-homologous protein (WAVE) heteropentameric regulatory complex (WRC)^13–17^. Cyfip2-mediated activation of the WRC requires it to bind with activated, GTP-bound Rac1^14,54,68^ to release WASF, which then triggers Arp2/3-mediated branched actin polymerization^14,54,68^. Our findings indicate that the Cyfip2-Rac1 interaction is required for the visually-driven dark flash, light flash, and prey capture behaviors while Cyfip2-FMRP/eIF4E interactions are dispensable (Figures 5-6). This contrasts with Cyfip2’s role in the acoustic startle circuit, in which loss of *cyfip2* causes hyperresponsiveness that requires both Cyfip2-Rac1 and Cyfip2-FMRP/eIF4E interactions to rescue^23^. Cyfip2 regulates the kinematics of both acoustic and visual startle responses, with *cyfip2* mutant responses having reduced turning angles^22,23^ (Figure 1). These weak motor responses are consistent with mutants’ reduced spontaneous locomotion^22^ and human DEE65 subjects’ hypotonia^55,56^. Cyfip2’s regulation of acoustic and visual response kinematics requires only Rac1 interactions and is independent of FMRP/eIF4E^23^ (Figure 6), and so these defects likely reflect a WRC-dependent role for Cyfip2 in spinal motor circuits. The molecular requirements for Cyfip2-mediated visual behaviors are also consistent with those needed for its role in retinotectal axon guidance. Cioni et al. demonstrated that Cyfip2 regulates pre-target segregation of dorsoventral RGC axons through binding with the WRC but not FMRP/eIF4e^12^. This observation aligns with previous reports that FMRP knockdown does not affect retinotectal topography in either wild-type or *cyfip2* deficient larvae, although broad translational inhibition does impair terminal RGC arborization in the optic tectum^11,69^.

The shared molecular requirements for Cyfip2-mediated RGC axon guidance and visual behavior suggest there may a functional connection between the two. Our data show that the optokinetic response (OKR) is intact in *cyfip2* mutants, aside from slight defects in eye abduction (Figure 2) that could be due to abducens nerve and/or lateral rectus muscle dysfunction. This is largely consistent with findings from mice in which *cyfip2* was conditionally knocked out in the retina. Electroretinograms showed no difference in photoreceptor or bipolar cell function, and multi-electrode recordings found only minor increases in ON RGC responses in *cyfip2* null retinas^43^. These mice were also capable of performing the OKR with only slight changes in the kinematics of eye movements^43^, which are more likely due to motor defects than to altered stimulus detection. That both zebrafish and mice lacking *cyfip2* have mostly intact OKRs suggests that Cyfip2 is not required for gross retinal function, despite defects in the lamination of some amacrine cells and RGCs in both species^11,43^ . And as the OKR is independent of the optic tectum in zebrafish^46^, this further implicates defects in the tectum as the most likely cause of *cyfip2* mutants’ deficient prey capture and dark flash/light flash responses. At 5 dpf Cyfip2 protein is strongly expressed in the tectum, along with the inner and outer plexiform layers of the retina, the olfactory bulb, and the lateral margins of the hindbrain^22^ . This doesn’t preclude function earlier as RGC axons are extending and making connections in the tectum, however, particularly as *cyfip2* mRNA is broadly expressed in the developing retina and CNS from 15-72 hpf^11^. Indeed, *cyfip2* is required cell-autonomously in RGCs for homotypic and heterotypic contact-triggered axon fasciculation and repulsion between dorsal and ventral RGC axons^11,12^, and axon-axon contact causes Cyfip2 to move into the growth cone where it regulates actin and filopodial dynamics^12^.

Our conditional rescue experiments further support a developmental role in RGC axon guidance for Cyfip2 in enabling behavioral responses to visual stimuli rather than in regulating acute circuit function. We found that expression of Cyfip2 between 30-50 hpf was sufficient to restore both dark flash/light flash responses and prey capture behavior in 5 dpf *cyfip2* mutants, while later expression had no effect (Figures 5-6). This critical window coincides with key steps in in the development of the visual system. Between 32-36 hpf RGC axons exit the eye and cross the optic chiasm, followed by their invasion of the optic tectum and 9 other arborization fields from 44-48 hpf^11,70–73^. Further topographic elaboration of RGC axon arbors continues through 72 hpf, after which the first visually-driven behaviors emerge^11,12,71,72^. Thus, the timing determined through our rescue experiments indicates that *cyfip2* is most critical for RGC axon guidance and sorting as they extend into the optic tectum. This correlates with the RGC axon defects observed in *cyfip2* mutants in which dorsonasal RGC axons take circuitous routes to their targets in the tectum by incorrectly entering the dorsal tectum through the dorsal rather than ventral branch of the optic tract^8,11^. Most axons eventually terminate correctly in the ventral tectum, however^8,11^, suggesting that *cyfip2* may also regulate synaptic connectivity and function in the tectum to enable proper visually-driven behaviors.

We used two approaches to measure how *cyfip2* regulates activity in the visual system. First, our brain-wide mapping of pERK staining showed that unstimulated activity in *cyfip2* mutants is substantially reduced in the tectum neuropil, along with multiple forebrain regions (Figure 3). This analysis also revealed a decrease in volume of the tectum neuropil, suggesting that RGC axon arborization and/or tectal neuron dendrite growth is reduced by the loss of *cyfip2,* which aligns with reports of reduced RGC arborization in the tectum^11^ and disrupted dendrite growth in the Mauthner neurons in the hindbrain^22^. With pan-neuronal calcium imaging we found that *cyfip2* does not regulate the probability of firing following a light stimulus for neurons in the pretectum, tectum neuropil, or cerebellum (Figure 4). The reduced peak change in GCaMP fluorescence in these neurons, however, suggests that when activated they fire fewer action potentials, likely limiting the downstream activation of hindbrain motor circuits. These experiments were limited by analyzing a single focal plane, however, and thus there may be more widespread defects in the activity of neurons in other layers of the tectum. Both of our approaches make clear, however, that activity in the tectum is reduced by loss of cyfip2, most likely as a result of diminished functional input from RGC axons.

## Conclusion

In this study we define how Cyfip2 functions as a critical mediator of vertebrate visual circuit assembly. We show that Cyfip2 acts through Rac1-dependent actin remodeling during a restricted developmental window between 30-50 hpf to establish functional visual circuits. These findings illuminate the mechanisms that contribute to visual deficits experienced by human subjects with mutations in CYFIP2. In future work, cell-type-specific manipulation of Cyfip2 signaling and analyses of synaptic connectivity and function between RGC and their targets will be instrumental in resolving how disrupted Cyfip2-mediated actin dynamics disrupt sensorimotor integration of visual stimuli in the tectum and hindbrain. Elucidating these mechanisms will clarify conserved principles of sensory circuit organization across species and inform the etiology of CYFIP2-associated neurodevelopmental disorders.

## Authors’ Contributions

*Project conceptualization*: Kimberly M. Charron and Kurt C. Marsden. *Methodology*: Kimberly M. Charron and Kurt C. Marsden. *Investigation*: Kimberly M. Charron, Sureni H. Sumathipala, D. Christopher Cole, Katya J. Frazier, Audrey L.B. Johnson, and Kurt C. Marsden. *Data curation*: Kimberly M. Charron and Melody B. Hancock. *Writing*: Kimberly M. Charron and Kurt C. Marsden. *Writing-review & editing*: Kimberly M. Charron and Kurt C. Marsden. *Supervision*: Kurt C. Marsden. *Funding acquisition*: Kurt C. Marsden.

## Supplementary Material

Supplementary Figure S1

Supplementary Figure S2

Supplementary Figure S3

Supplementary Figure S4

Supplementary Figure S5

Supplementary Figure S6

Supplementary Figure S7

Supplementary Figure S8

Supplementary Figure S9

Supplementary Video S1

Supplementary Video S2

Supplementary Video S3

Supplementary Video S4

Supplementary Video S5

Supplementary Video S6

Supplementary Table S1

## Acknowledgements

We would like to thank Dr. Jessica Nelson (University of Colorado, Anschutz Medical Campus, Department of Cell and Developmental Biology) and Dr. Summer Thyme (UMass Chan Medical School, Department of Biochemistry and Molecular Biotechnology) for their assistance with MAP-mapping analysis. We would also like to thank Dr. John Hageter and Dr. Eric Horstick (West Virginia University, Department of Biology) for the development and troubleshooting of the MCA pipeline.

## Reference

1. Kaas JH. Topographic maps are fundamental to sensory processing. Brain Res Bull. 1997;44(2):107–112.

2. Suzuki DG, Pérez-Fernández J, Wibble T, Kardamakis AA, Grillner S. The role of the optic tectum for visually evoked orienting and evasive movements. Proc Natl Acad Sci U S A. 2019;116(30):15272–15281.

3. Isa T, Marquez-Legorreta E, Grillner S, Scott EK. The tectum/superior colliculus as the vertebrate solution for spatial sensory integration and action. Curr Biol. 2021;31(11):R741–R762.

4. Robles E, Laurell E, Baier H. The retinal projectome reveals brain-area-specific visual representations generated by ganglion cell diversity. Curr Biol. 2014;24(18):2085–2096.

5. Novales Flamarique I, Wachowiak M. Functional segregation of retinal ganglion cell projections to the optic tectum of rainbow trout. J Neurophysiol. 2015;114(5):2703–2717.

6. Kukreja S, Gautam P, Saxena R, et al. Identification of novel candidate regulators of retinotectal map formation through transcriptional profiling of the chick optic tectum. J Comp Neurol. 2017;525(3):459–477.

7. Xu NJ, Henkemeyer M. Ephrin reverse signaling in axon guidance and synaptogenesis. Semin Cell Dev Biol. 2012;23(1):58–64.

8. Trowe T, Klostermann S, Baier H, et al. Mutations disrupting the ordering and topographic mapping of axons in the retinotectal projection of the zebrafish, Danio rerio. Development. 1996;123:439–450.

9. Karlstrom RO, Trowe T, Klostermann S, et al. Zebrafish mutations affecting retinotectal axon pathfinding. Development. 1996;123:427–438.

10. Baier H, Klostermann S, Trowe T, Karlstrom RO, Nüsslein-Volhard C, Bonhoeffer F. Genetic dissection of the retinotectal projection. Development. 1996;123:415–425.

11. Pittman AJ, Gaynes JA, Chien CB. Nev (cyfip2) is required for retinal lamination and axon guidance in the zebrafish retinotectal system. Dev Biol. 2010;344(2):784–794.

12. Cioni JM, Wong HHW, Bressan D, Kodama L, Harris WA, Holt CE. Axon-Axon Interactions Regulate Topographic Optic Tract Sorting via CYFIP2-Dependent WAVE Complex Function. Neuron. 2018;97(5):1078–1093.e6.

13. Schenck A, Bardoni B, Moro A, Bagni C, Mandel JL. A highly conserved protein family interacting with the fragile X mental retardation protein (FMRP) and displaying selective interactions with FMRP-related proteins FXR1P and FXR2P. Proc Natl Acad Sci U S A. 2001;98(15):8844–8849.

14. Chen Z, Borek D, Padrick SB, et al. Structure and control of the actin regulatory WAVE complex. Nature. 2010;468(7323):533–538.

15. Cory GOC, Ridley AJ. Cell motility: braking WAVEs. Nature. 2002;418(6899):732–733.

16. Eden S, Rohatgi R, Podtelejnikov AV, Mann M, Kirschner MW. Mechanism of regulation of WAVE1-induced actin nucleation by Rac1 and Nck. Nature. 2002;418(6899):790–793.

17. Chen B, Brinkmann K, Chen Z, et al. The WAVE regulatory complex links diverse receptors to the actin cytoskeleton. Cell. 2014;156(1-2):195–207.

18. Takenawa T, Suetsugu S. The WASP-WAVE protein network: connecting the membrane to the cytoskeleton. Nat Rev Mol Cell Biol. 2007;8(1):37–48.

19. Derivery E, Lombard B, Loew D, Gautreau A. The Wave complex is intrinsically inactive. Cell Motil Cytoskeleton. 2009;66(10):777–790.

20. Napoli I, Mercaldo V, Boyl PP, et al. The fragile X syndrome protein represses activity-dependent translation through CYFIP1, a new 4E-BP. Cell. 2008;134(6):1042–1054.

21. De Rubeis S, Pasciuto E, Li KW, et al. CYFIP1 coordinates mRNA translation and cytoskeleton remodeling to ensure proper dendritic spine formation. Neuron. 2013;79(6):1169–1182.

22. Marsden KC, Jain RA, Wolman MA, et al. A Cyfip2-Dependent Excitatory Interneuron Pathway Establishes the Innate Startle Threshold. Cell Rep. 2018;23(3):878–887.

23. Deslauriers JC, Ghotkar RP, Russ LA, et al. Cyfip2 controls the acoustic startle threshold through FMRP, actin polymerization, and GABAB receptor function. bioRxiv. Published online December 22, 2023. doi:10.1101/2023.12.22.573054

24. Randlett O, Wee CL, Naumann EA, et al. Whole-brain activity mapping onto a zebrafish brain atlas. Nat Methods. 2015;12(11):1039–1046.

25. Capps MES, Moyer AJ, Conklin CL, et al. Disrupted diencephalon development and neuropeptidergic pathways in zebrafish with autism-risk mutations. Proc Natl Acad Sci U S A. 2025;122(23):e2402557122.

26. Rohlfing T, Maurer CR. Nonrigid image registration in shared-memory multiprocessor environments with application to brains, breasts, and bees. IEEE Trans Inf Technol Biomed. 2003;7(1):16–25.

27. Jefferis GSXE, Potter CJ, Chan AM, et al. Comprehensive maps of Drosophila higher olfactory centers: spatially segregated fruit and pheromone representation. Cell. 2007;128(6):1187–1203.

28. Gu Z. Complex heatmap visualization. Imeta. 2022;1(3):e43.

29. Gu Z, Eils R, Schlesner M. Complex heatmaps reveal patterns and correlations in multidimensional genomic data. Bioinformatics. 2016;32(18):2847–2849.

30. Hageter J, DelGaudio A, Leathery M, et al. MCA: A Multicellular analysis Calcium Imaging toolbox for ImageJ. *bioRxivorg*. Published online August 23, 2025. doi:10.1101/2025.08.19.671108

31. Vincent L, Soille P. Watersheds in digital spaces: an efficient algorithm based on immersion simulations. IEEE Trans Pattern Anal Mach Intell. 1991;13(6):583–598.

32. Burgess HA, Granato M. Sensorimotor gating in larval zebrafish. Journal of Neuroscience. 2007;27(18):4984–4994.

33. Wolman MA, Jain RA, Liss L, Granato M. Chemical modulation of memory formation in larval zebrafish. Proc Natl Acad Sci U S A. 2011;108(37):15468–15473.

34. Burgess HA, Granato M. Modulation of locomotor activity in larval zebrafish during light adaptation. J Exp Biol. 2007;210(Pt 14):2526–2539.

35. Rodwell V, Birchall A, Yoon HJ, Kuht HJ, Norton WHJ, Thomas MG. A novel portable flip-phone based visual behaviour assay for zebrafish. Sci Rep. 2024;14(1):236.

36. Scheetz SD, Shao E, Zhou Y, Cario CL, Bai Q, Burton EA. An open-source method to analyze optokinetic reflex responses in larval zebrafish. J Neurosci Methods. 2018;293:329–337.

37. Patterson BW, Abraham AO, MacIver MA, McLean DL. Visually guided gradation of prey capture movements in larval zebrafish. J Exp Biol. 2013;216(Pt 16):3071–3083.

38. Gahtan E, Tanger P, Baier H. Visual prey capture in larval zebrafish is controlled by identified reticulospinal neurons downstream of the tectum. J Neurosci. 2005;25(40):9294–9303.

39. Bianco IH, Kampff AR, Engert F. Prey capture behavior evoked by simple visual stimuli in larval zebrafish. Front Syst Neurosci. 2011;5:101.

40. Randlett O, Haesemeyer M, Forkin G, et al. Distributed Plasticity Drives Visual Habituation Learning in Larval Zebrafish. Curr Biol. 2019;29(8):1337–1345.e4.

41. Hodorovich DR, Lindsley PM, Berry AA, Burton DF, Marsden KC. Morphological and sensorimotor phenotypes in a zebrafish CHARGE syndrome model are domain-dependent. Genes Brain Behav. 2023;22(3):e12839.

42. Martin RM, Bereman MS, Marsden KC. BMAA and MCLR interact to modulate behavior and exacerbate molecular changes related to neurodegeneration in larval zebrafish. Toxicol Sci. 2021;179(2):251–261.

43. Chaya T, Ishikane H, Varner LR, et al. Deficiency of the neurodevelopmental disorder-associated gene Cyfip2 alters the retinal ganglion cell properties and visual acuity. Hum Mol Genet. 2022;31(4):535–547.

44. Schweigart G, Mergner T, Evdokimidis I, Morand S, Becker W. Gaze stabilization by optokinetic reflex (OKR) and vestibulo-ocular reflex (VOR) during active head rotation in man. Vision Res. 1997;37(12):1643–1652.

45. Matsuda K, Kubo F. Circuit organization underlying optic flow processing in zebrafish. Front Neural Circuits. 2021;15:709048.

46. Roeser T, Baier H. Visuomotor behaviors in larval zebrafish after GFP-guided laser ablation of the optic tectum. J Neurosci. 2003;23(9):3726–3734.

47. Mustari MJ, Ono S. Optokinetic Eye Movements. In: *Encyclopedia of Neuroscience*. Elsevier; 2009:285-293.

48. Scully C. Neurology. In: *Scully’s Medical Problems in Dentistry*. Elsevier; 2014:345-392.

49. Duchemin A, Privat M, Sumbre G. Fourier motion processing in the optic tectum and pretectum of the zebrafish larva. Front Neural Circuits. 2021;15:814128.

50. Knorr AG, Gravot CM, Glasauer S, Straka H. Image motion with color contrast suffices to elicit an optokinetic reflex in Xenopus laevis tadpoles. Sci Rep. 2021;11(1):8445.

51. Stringer C, Wang T, Michaelos M, Pachitariu M. Cellpose: a generalist algorithm for cellular segmentation. Nat Methods. 2021;18(1):100–106.

52. Spranger S, Rommel B, Jauch A, Bodammer R, Mehl B, Bullerdiek J. Interstitial deletion of 5q33.3q35.1 in a girl with mild mental retardation. Am J Med Genet. 2000;93(2):107–109.

53. Lee JH, Kim HJ, Yoon JM, et al. Interstitial deletion of 5q33.3q35.1 in a boy with severe mental retardation. Korean J Pediatr. 2016;59(Suppl 1):S19–S24.

54. Nakashima M, Kato M, Aoto K, et al. De novo hotspot variants in CYFIP2 cause early-onset epileptic encephalopathy. Ann Neurol. 2018;83(4):794–806.

55. Zweier M, Begemann A, McWalter K, et al. Spatially clustering de novo variants in CYFIP2, encoding the cytoplasmic FMRP interacting protein 2, cause intellectual disability and seizures. Eur J Hum Genet. 2019;27(5):747–759.

56. Begemann A, Sticht H, Begtrup A, et al. New insights into the clinical and molecular spectrum of the novel CYFIP2-related neurodevelopmental disorder and impairment of the WRC-mediated actin dynamics. Genet Med. 2021;23(3):543–554.

57. Deslauriers J. Cyfip2 Controls the Acoustic Startle Threshold and Visually Mediated Behaviors. North Carolina State University ; 2023.

58. Schenck A, Bardoni B, Langmann C, Harden N, Mandel JL, Giangrande A. CYFIP/Sra-1 controls neuronal connectivity in Drosophila and links the Rac1 GTPase pathway to the fragile X protein. Neuron. 2003;38(6):887–898.

59. Saller E, Tom E, Brunori M, et al. Increased apoptosis induction by 121F mutant p53. EMBO J. 1999;18(16):4424–4437.

60. Freeman J, Schmidt S, Scharer E, Iggo R. Mutation of conserved domain II alters the sequence specificity of DNA binding by the p53 protein. EMBO J. 1994;13(22):5393–5400.

61. Schenck A, Qurashi A, Carrera P, et al. WAVE/SCAR, a multifunctional complex coordinating different aspects of neuronal connectivity. Dev Biol. 2004;274(2):260–270.

62. Borla MA, Palecek B, Budick S, O’Malley DM. Prey capture by larval zebrafish: evidence for fine axial motor control. Brain Behav Evol. 2002;60(4):207–229.

63. Fernandes AM, Fero K, Arrenberg AB, Bergeron SA, Driever W, Burgess HA. Deep brain photoreceptors control light-seeking behavior in zebrafish larvae. Curr Biol. 2012;22(21):2042–2047.

64. Lamiré LA, Haesemeyer M, Engert F, Granato M, Randlett O. Functional and pharmacological analyses of visual habituation learning in larval zebrafish. Elife. 2023;12. doi:10.7554/eLife.84926

65. Huang KH, Ahrens MB, Dunn TW, Engert F. Spinal projection neurons control turning behaviors in zebrafish. Curr Biol. 2013;23(16):1566–1573.

66. Orger MB, Kampff AR, Severi KE, Bollmann JH, Engert F. Control of visually guided behavior by distinct populations of spinal projection neurons. Nat Neurosci. 2008;11(3):327–333.

67. Easter SS Jr, Nicola GN. The development of vision in the zebrafish (Danio rerio). Dev Biol. 1996;180(2):646–663.

68. Chen B, Chou HT, Brautigam CA, et al. Rac1 GTPase activates the WAVE regulatory complex through two distinct binding sites. Elife. 2017;6. doi:10.7554/eLife.29795

69. Wong HHW, Lin JQ, Ströhl F, et al. RNA Docking and Local Translation Regulate Site-Specific Axon Remodeling In Vivo. Neuron. 2017;95(4):852–868.e8.

70. Hu M, Easter SS. Retinal neurogenesis: the formation of the initial central patch of postmitotic cells. Dev Biol. 1999;207(2):309–321.

71. Stuermer CA. Retinotopic organization of the developing retinotectal projection in the zebrafish embryo. J Neurosci. 1988;8(12):4513–4530.

72. Burrill JD, Easter SS Jr. Development of the retinofugal projections in the embryonic and larval zebrafish (Brachydanio rerio). J Comp Neurol. 1994;346(4):583–600.

73. Bollmann JH. The Zebrafish Visual System: From Circuits to Behavior. Annu Rev Vis Sci. 2019;5:269–293.

