## Supplementary Fig. S1 for "Cyfip2 mediates sensorimotor integration of visual input through Rac1-dependent actin remodeling"

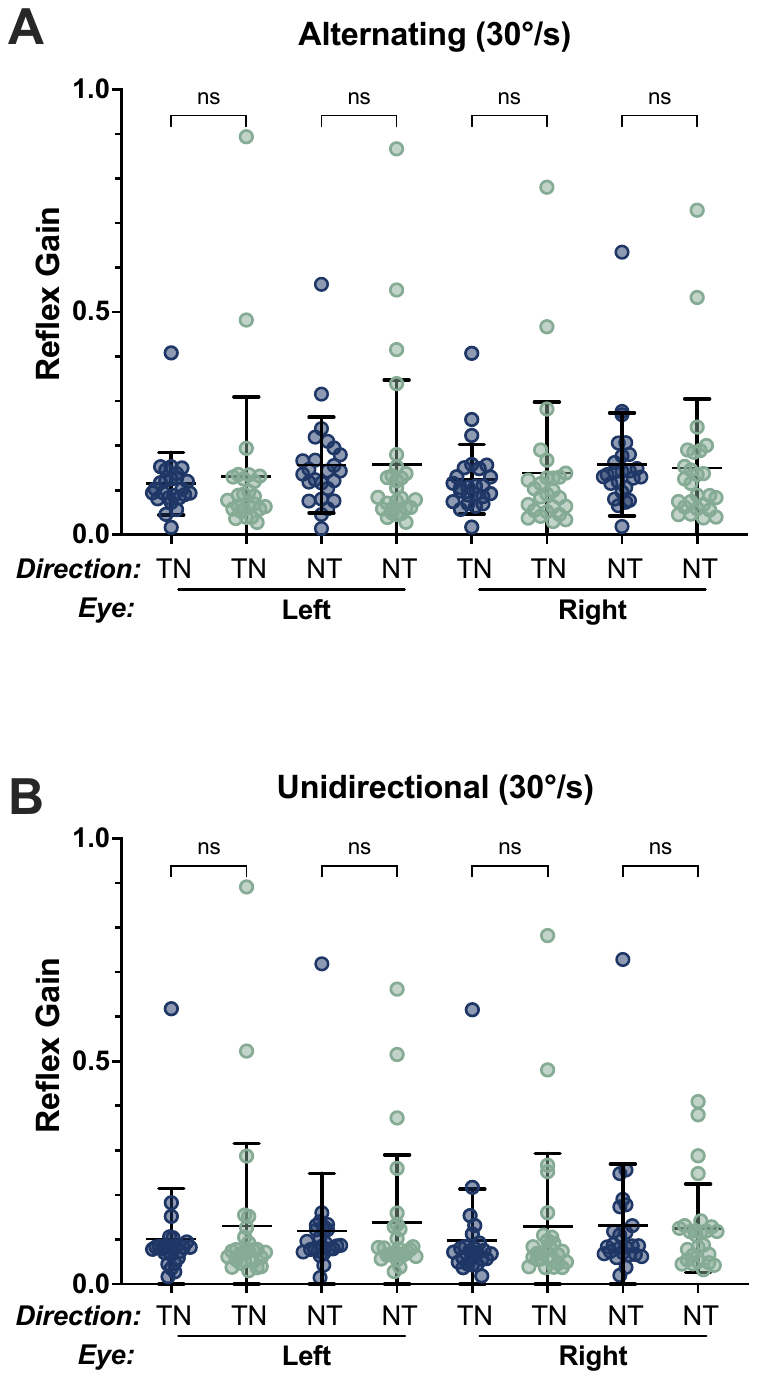

**Supplementary Figure S1: *cyfip2* mutants have subtle defects in the Optokinetic Response.** Quantification of optokinetic response to 99% contrast sinusoidal grating moving in either alternating or unidirectional pattern at a velocity 30°/s. Reflex gain of left or right eye in nasotemporal (NT) or temporonasal (TN) direction during alternating **A)** and unidirectional **B)** phase of 30°/s sinusoidal grating of *cyfip2^+/^* (n=25) and *cyfip2^-/-^* (n=26) larvae (mean ± SD). Comparisons made using 1-way ANOVA with Šídák’s multiple comparisons test.

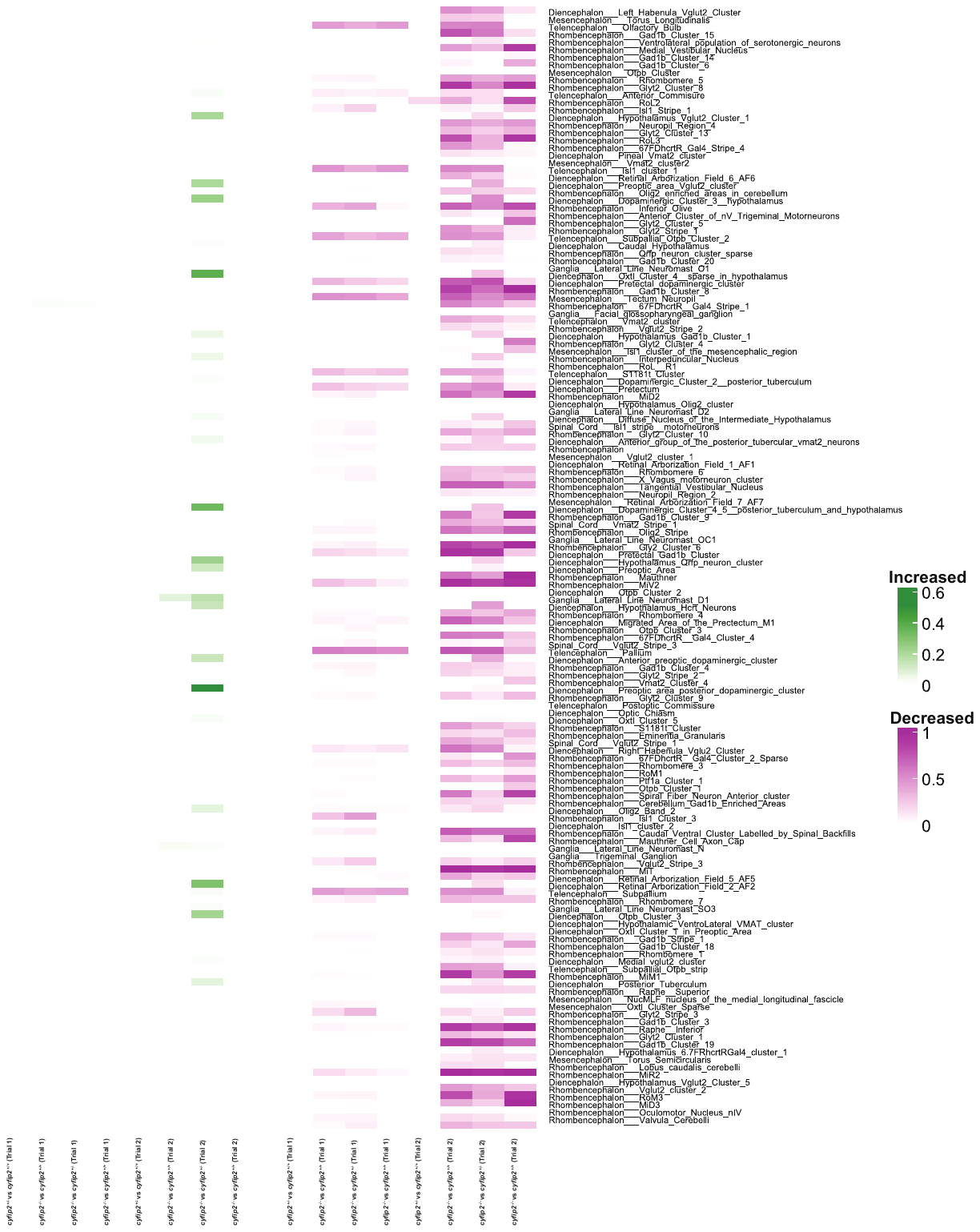

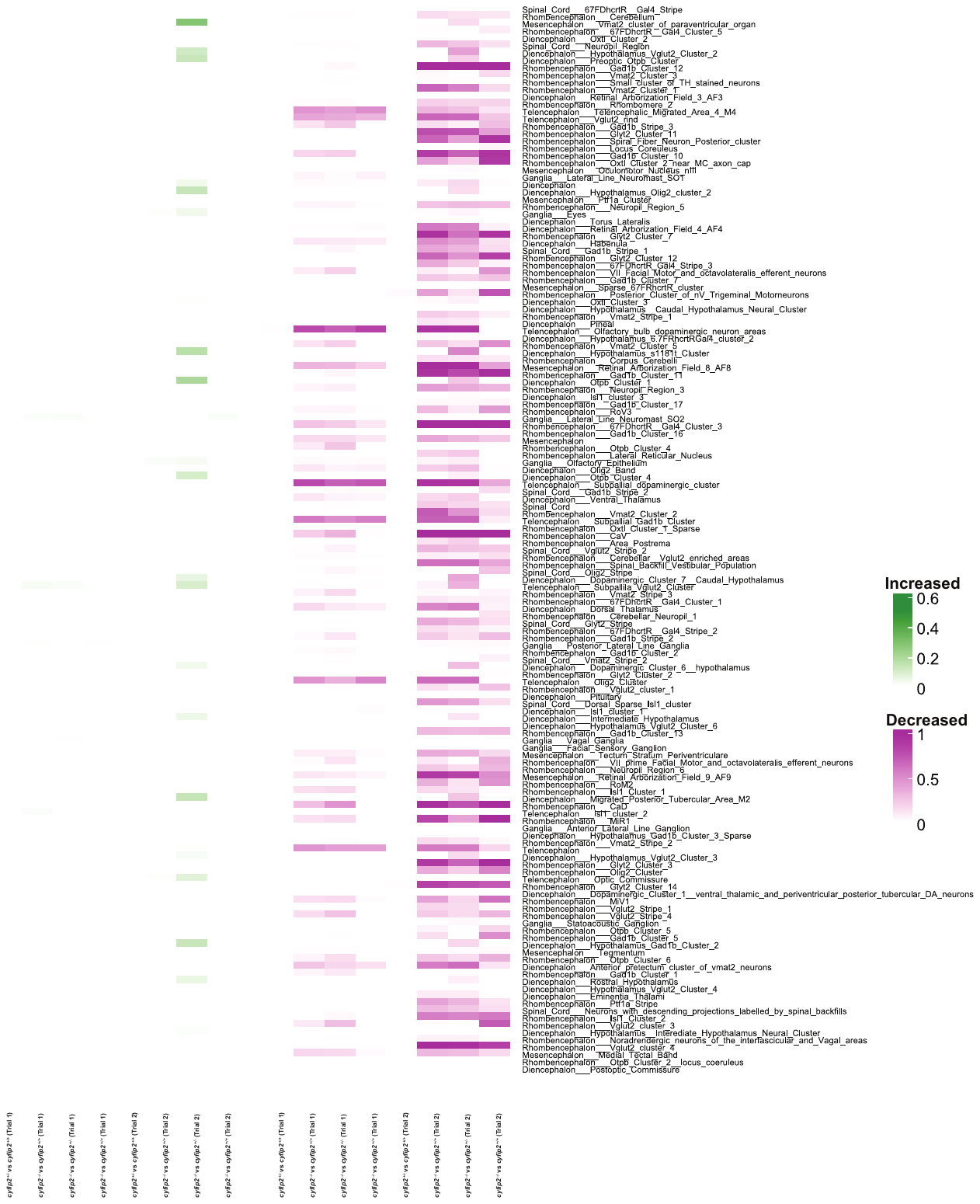

**Supplementary Figure S2: *cyfip2* mutants display both a decrease in both activity and brain volume within the optic tectum.** Heatmap of changes in activity across all 294 brain regions and genotype comparisons where decreases in activity are shown in magenta and increases in activity are shown in green.

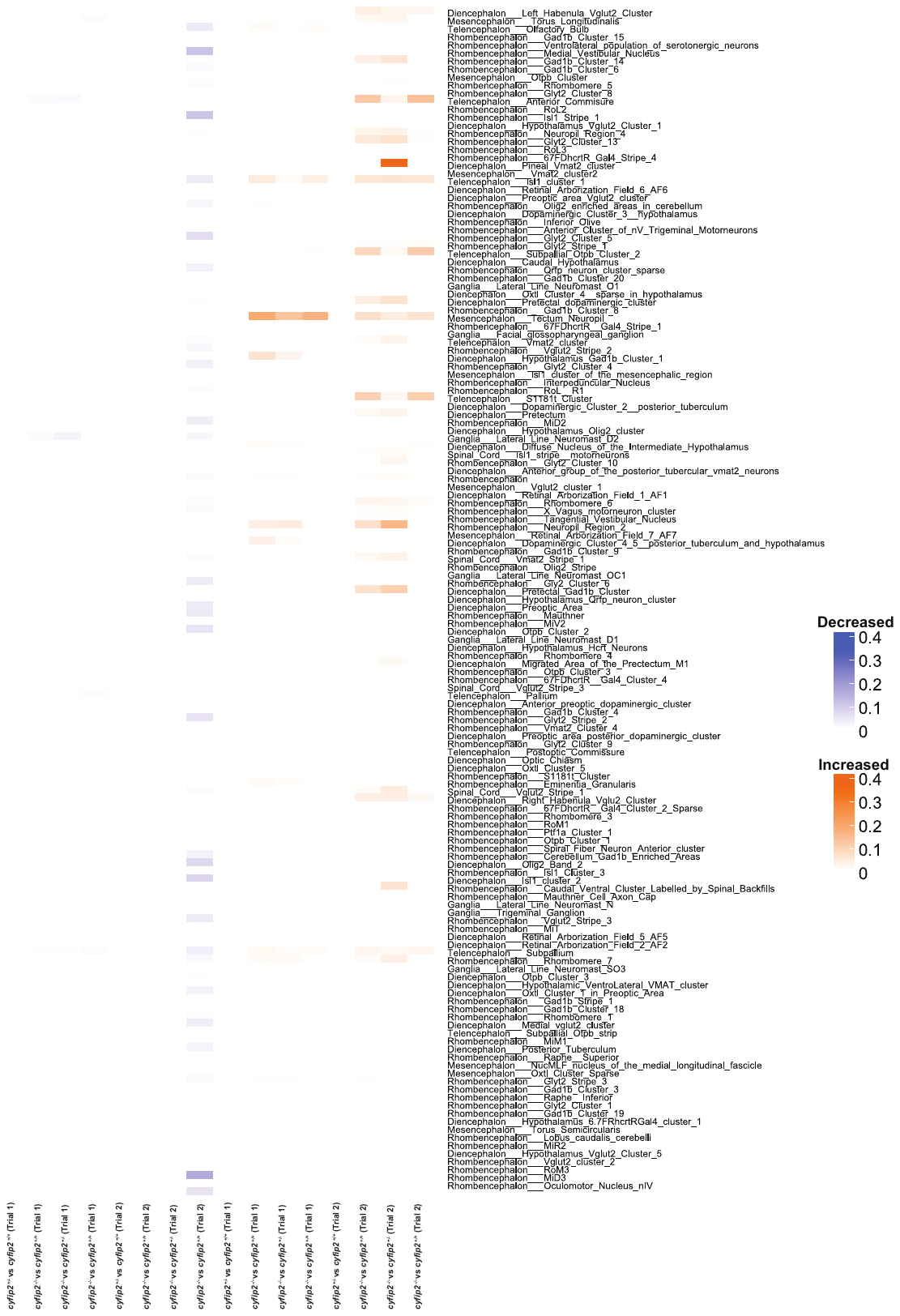

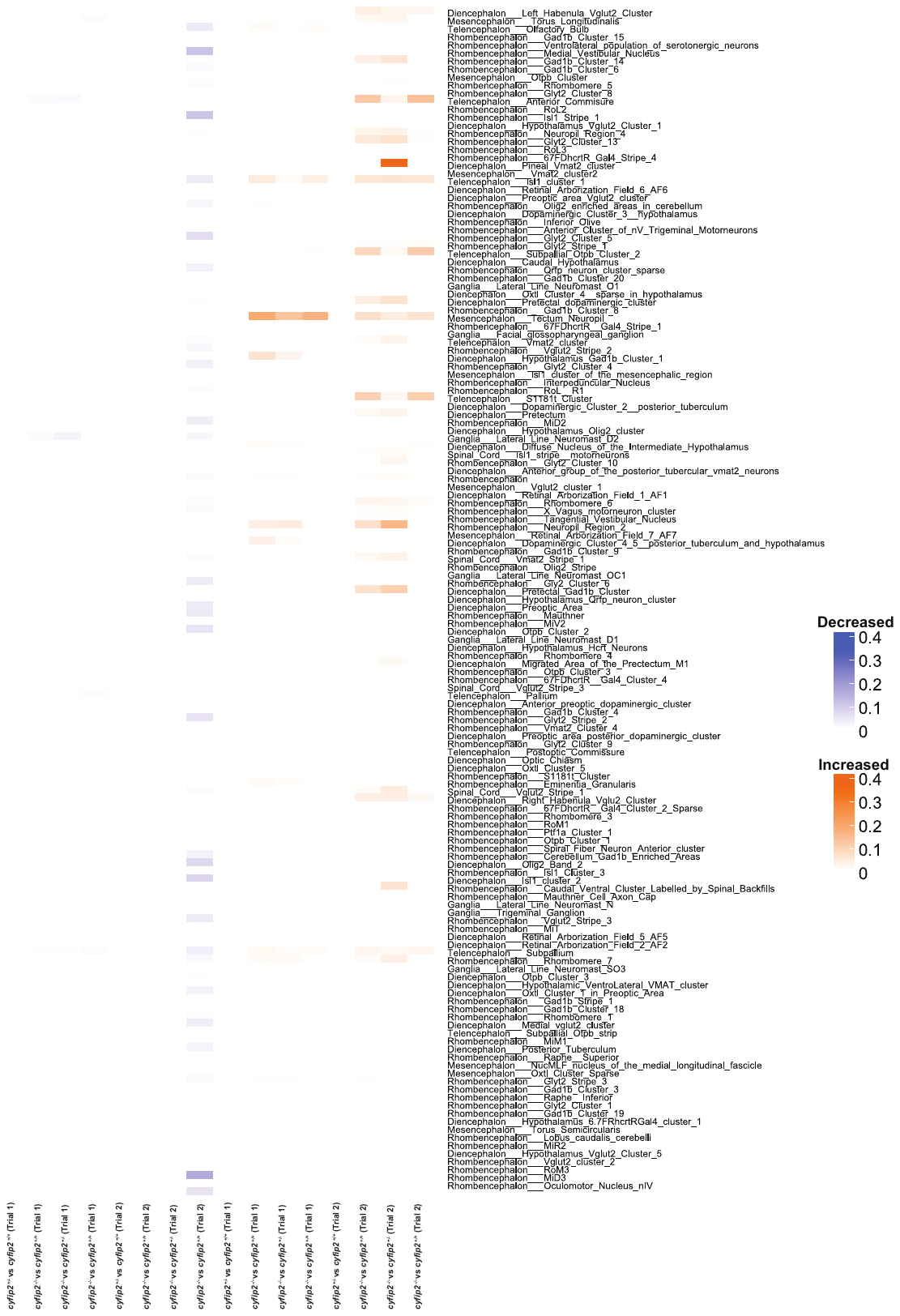

**Supplementary Figure S3: *cyfip2* mutants display both a decrease in both activity and brain volume within the optic tectum.** Heatmap of changes in morphometry across all 294 brain regions and genotype comparisons where decreases in morphometry are shown in blue and increases in morphometry are shown in orange.

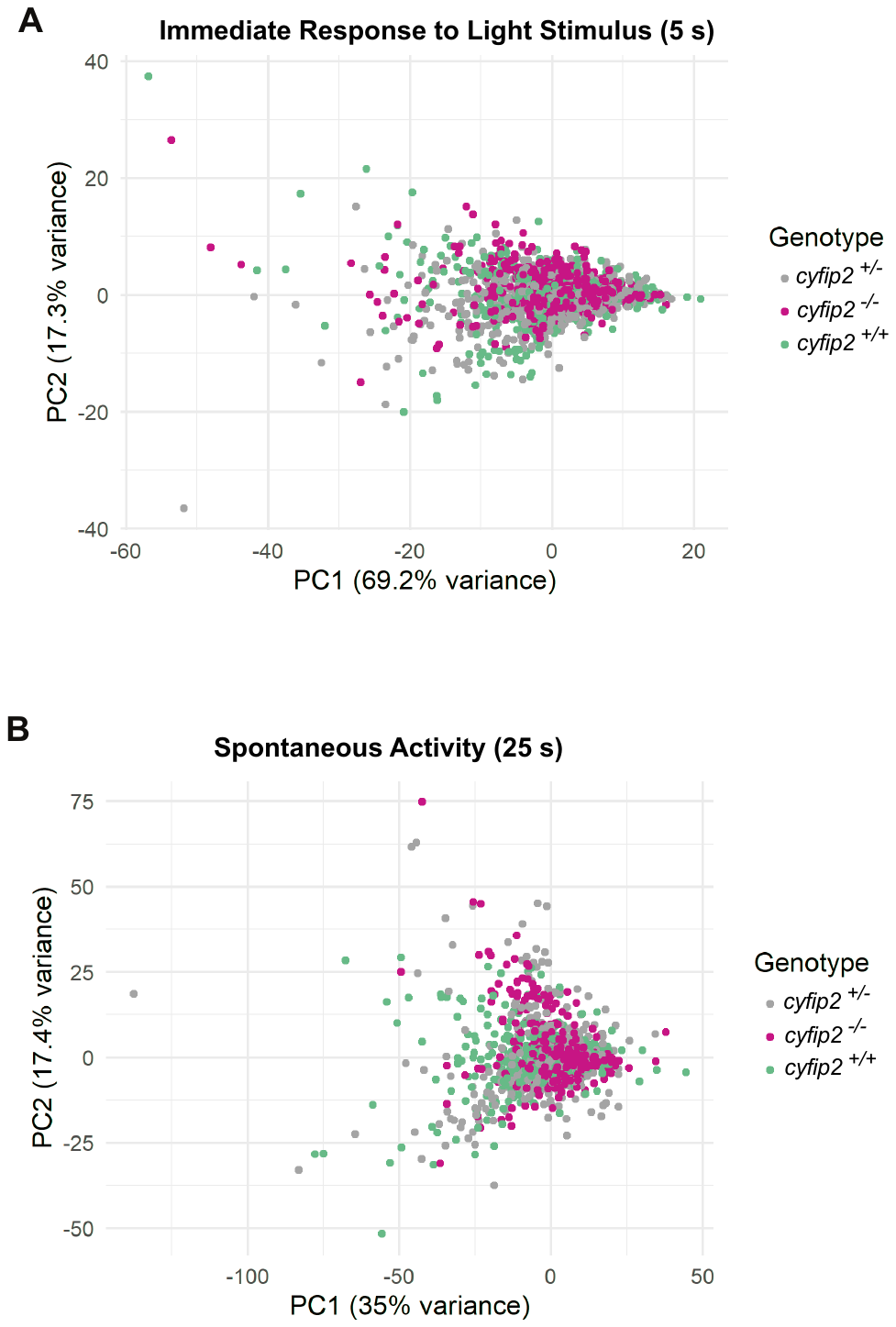

**Supplementary Figure S4: Absence of Cyfip2 leads to dampened neuronal signaling following light stimulus**. Principal component analysis (PCA) of **A)** the immediate response to the light stimulus and **B)** spontaneous activity following light stimulus.

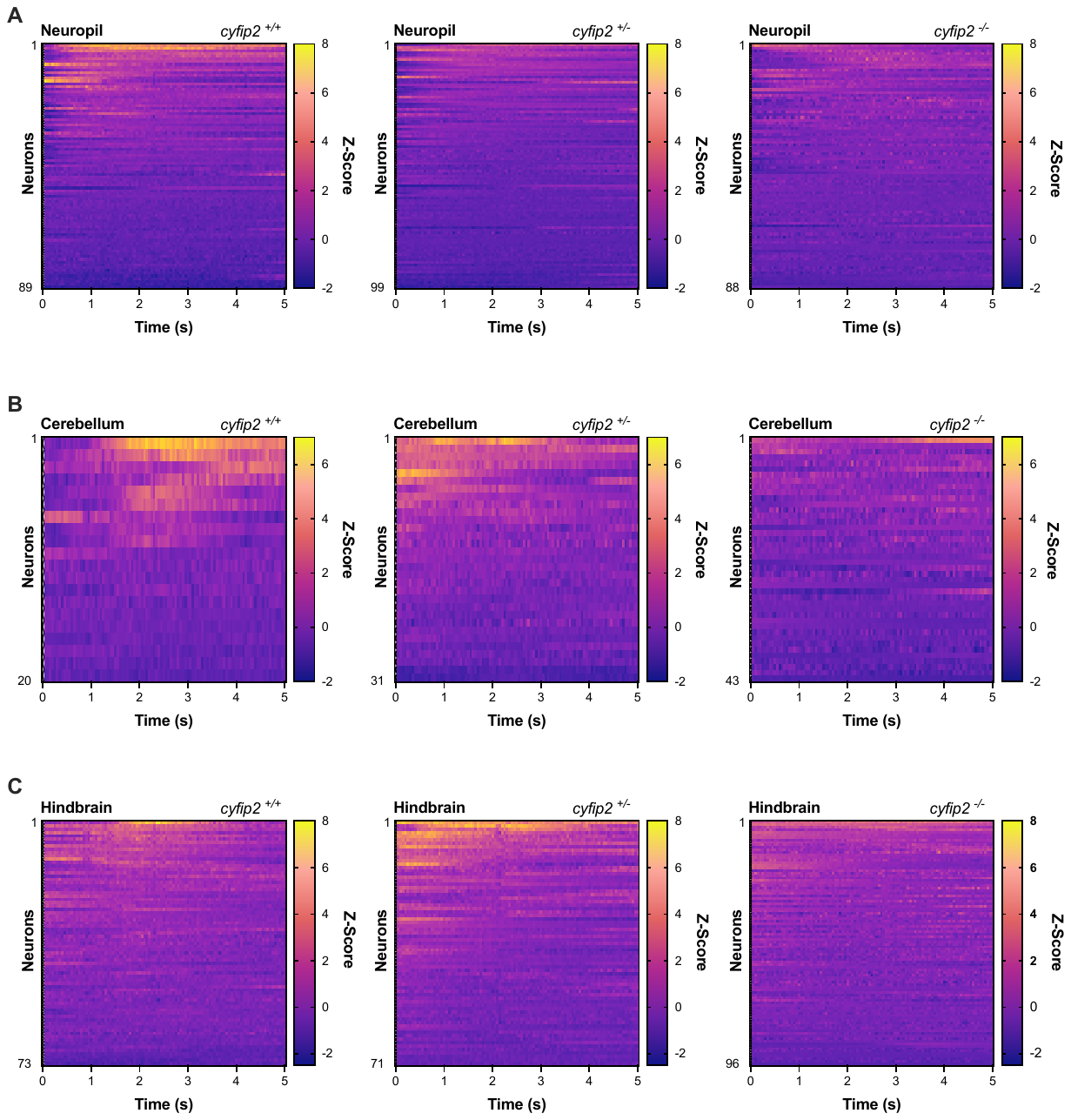

**Supplementary Figure S5: Absence of Cyfip2 leads to dampened neuronal signaling following light stimulus**. Normalized heatmaps depicting neural activity in the Neuropil **A)** the Cerebellum **B)** and the Hindbrain **C)** in the first 5 seconds following a blue light (488nm) stimulus of cyfip2^+/+^, cyfip2^+/-^ and cyfip2^-/-^.
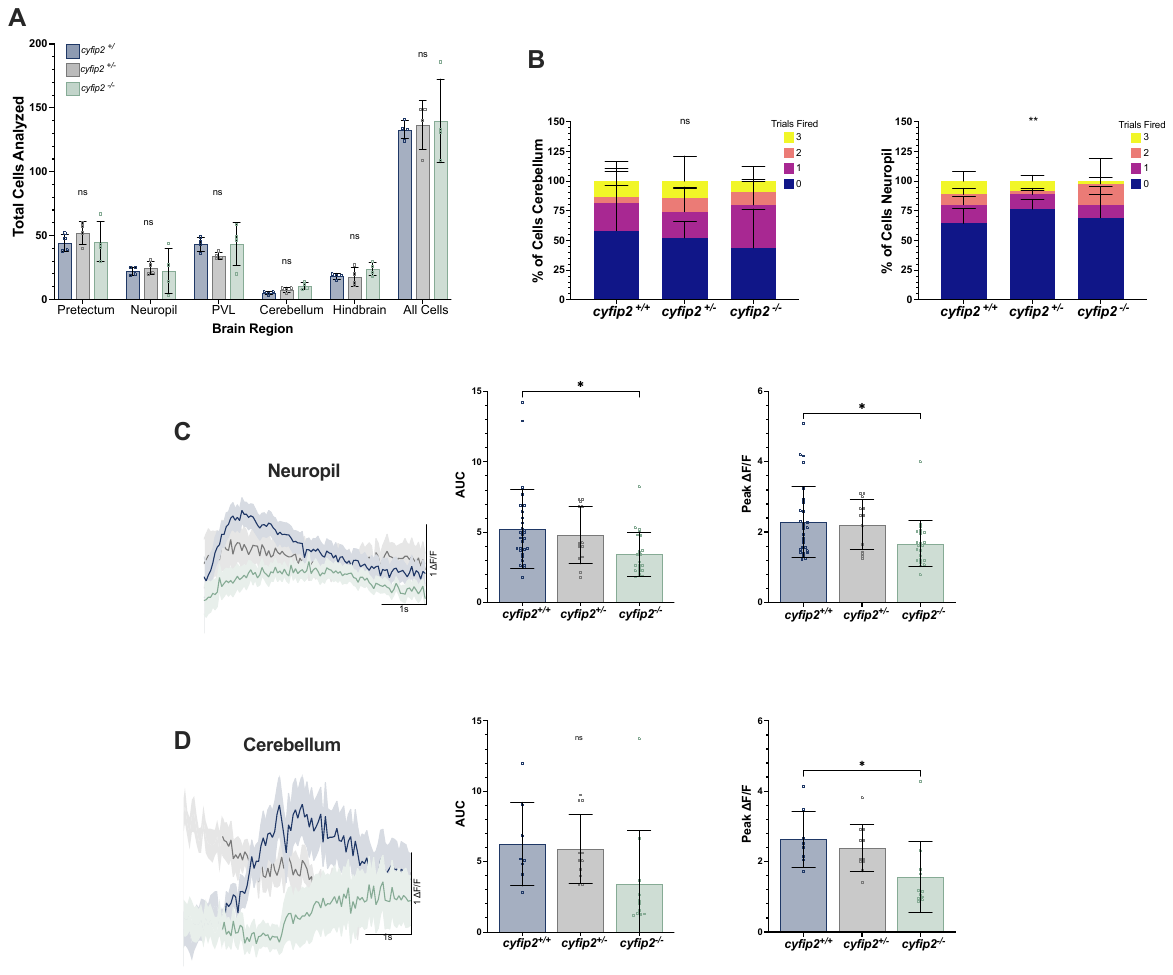

**Supplementary Figure S6: Absence of Cyfip2 leads to dampened neuronal signaling following light stimulus**. **A)** Total number of analyzed cells in the Pretectum, Neuropil, Periventricular Layer (PVL), Cerebellum, and Hindbrain of cyfip2^+/+^, cyfip2^+/-^ and cyfip2^-/-^ larvae. **B)** The percentage of neurons that fire (Z score >1.5) over the course of three successive trials, 3 times (yellow), 2 times (orange), 1 time (magenta), and 0 times (dark purple), in the Cerebellum and Neuropil across all three genotypes. Chi square statistical analysis used to determine significance. **C-D)** The average ∆F/F trace of each neuron that fired, the area under the curve (AUC) of the ∆F/F trace or each active neuron, and the peak ∆F/F value for cyfip2^+/+^ (blue), cyfip2^+/-^ (grey) and cyfip2^-/-^ in all cells, the Neuropil and Cerebellum. Data are presented as mean ± SEM for the average ∆F/F traces, and mean ± SD for all other data. Comparisons made using 1-way ANOVA with Tukey’s multiple comparisons test, * p < 0.05, **p < 0.01, ***p < 0.001, ****p < 0.0001.

**Supplementary Figure S7: Absence of Cyfip2 leads to dampened neuronal signaling following light stimulus**. Normalized heatmaps depicting spontaneous neural activity in all cells **A)** the Pretectum **B)** the Neuropil **C)** the Periventricular layer **D)** the Cerebellum **E)** and the Hindbrain **F)** from 5 to 30 seconds following a blue light (488nm) stimulus of cyfip2^+/+^, cyfip2^+/-^ and cyfip2^-/-^.

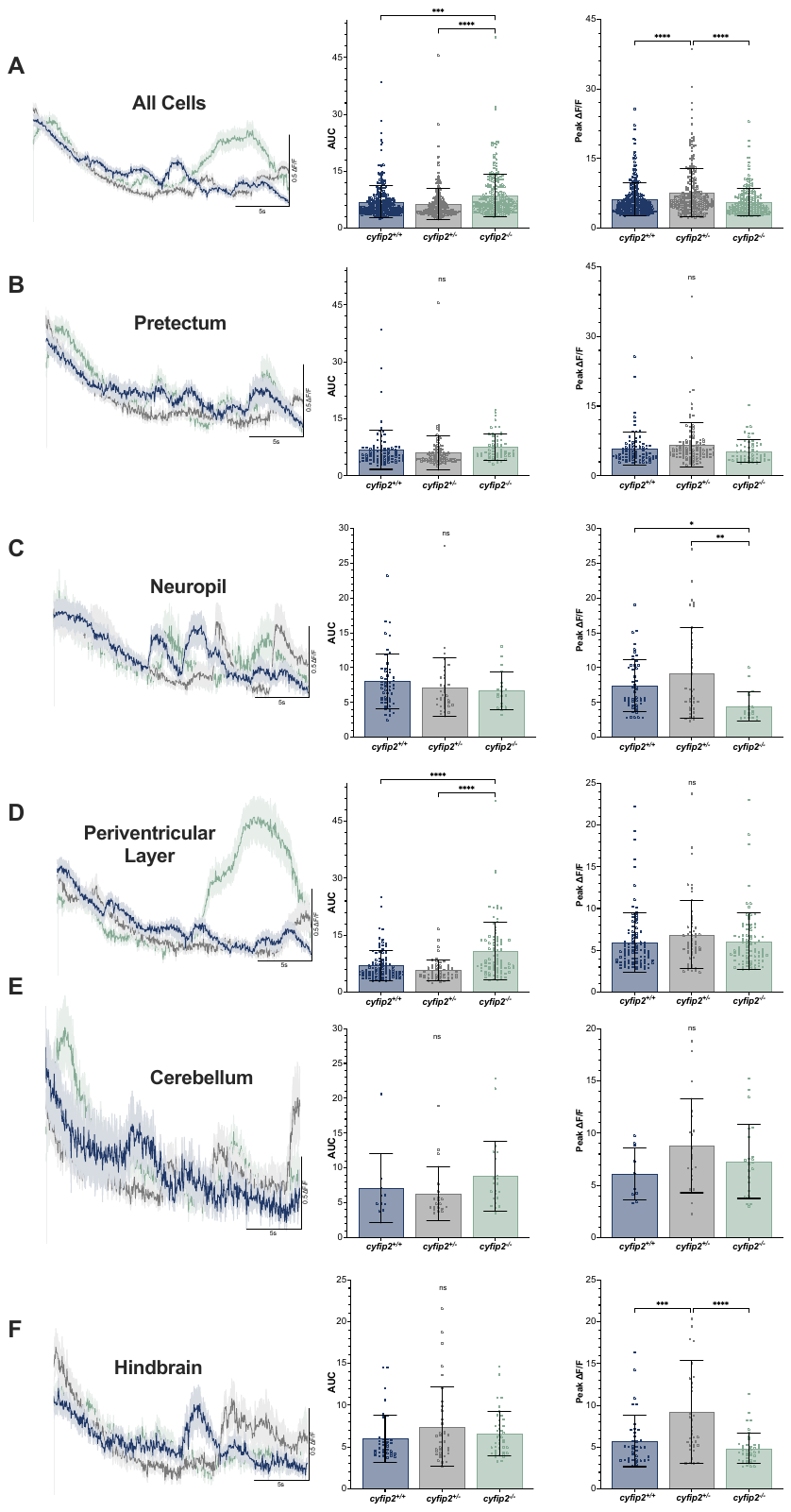

**Supplementary Figure S8: Absence of Cyfip2 leads to dampened neuronal signaling following light stimulus**. **E-H)** The average ∆F/F trace of each neuron that fired, the area under the curve (AUC) of the ∆F/F trace or each active neuron, and the peak ∆F/F value for cyfip2^+/+^ (blue), cyfip2^+/-^ (grey) and cyfip2^-/-^ in all cells, the Pretectum, Neuropil, Periventricular Layer, Cerebellum, and the Hindbrain from 5 to 30 seconds following a blue light (488nm) stimulus of cyfip2^+/+^, cyfip2^+/-^ and cyfip2^-/-^. Data are presented as mean ± SEM for the average ∆F/F traces, and mean ± SD for all other data. Comparisons made using 1-way ANOVA with Tukey’s multiple comparisons test, * p < 0.05, **p < 0.01, ***p < 0.001, ****p < 0.0001.

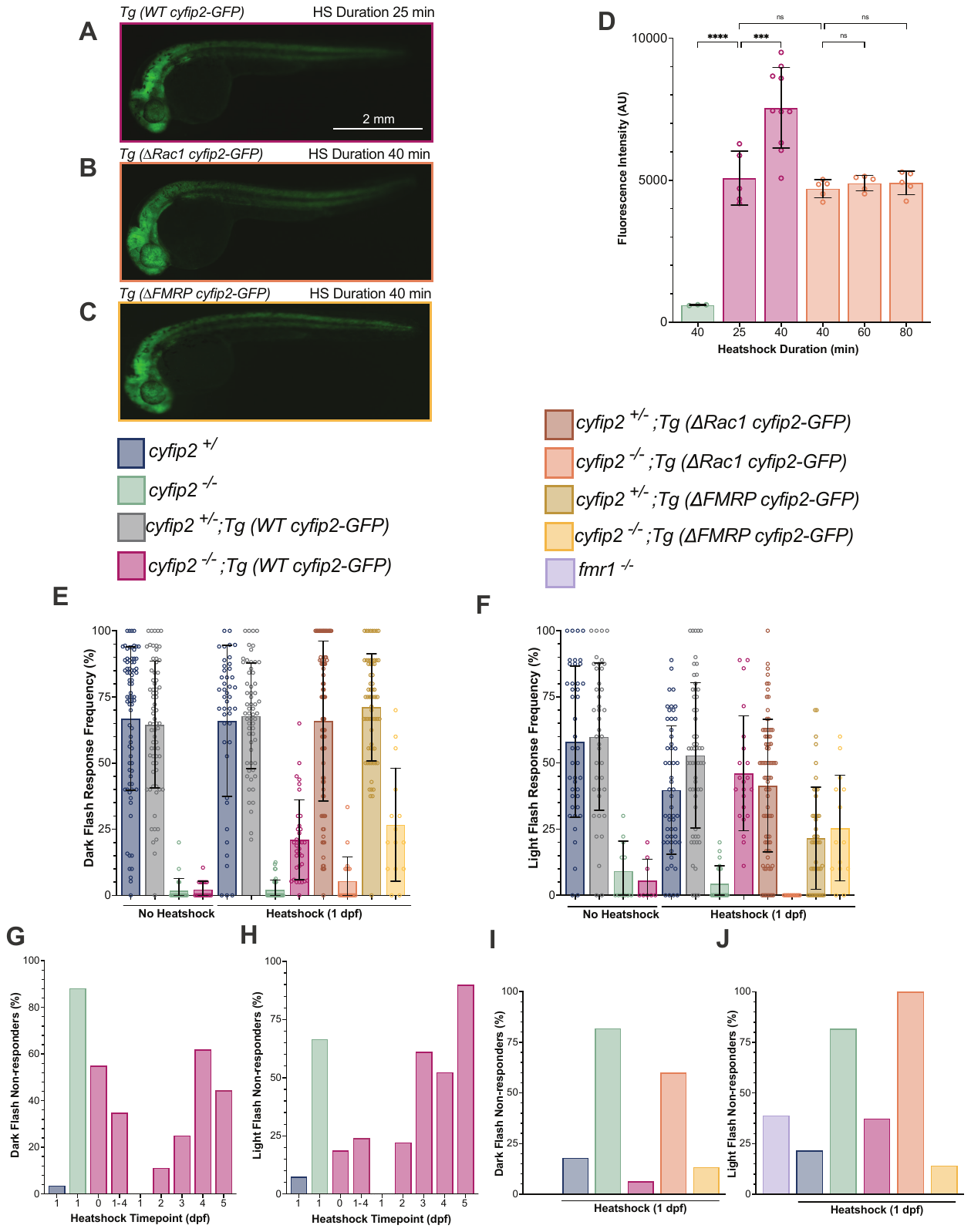

**Supplementary Figure S9: *cyfip2* developmentally regulates visual behavior through Rac1-dependent actin remodeling**. **A-C).** Representative fluorescence images of *cyfip2^-/-^;Tg(WT cyfip2-GFP)* (magenta), *cyfip2^-/-^;Tg(∆Rac1 cyfip2-GFP)* (orange), and *cyfip2^-/-^;Tg(∆FMRP cyfip2-GFP)* (yellow bar) at 34 hpf after 38°C heatshock at 30 hpf. **D)** Quantified Fluorescence intensity by heatshock duration of *cyfip2^-/-^;Tg(WT cyfip2-GFP)*, *cyfip2^-/-^;Tg(∆Rac1 cyfip2-GFP)* (orange), and *cyfip2^-/-^* (green). **E)** Response frequency to 10 dark or **F)** light flash stimuli at 5dpf of *cyfip2^+/^, cyfip2^-/-^, cyfip2^+/^;Tg(WT cyfip2-GFP), cyfip2^-/-^;Tg(WT cyfip2-GFP), cyfip2^+/-^;Tg(∆Rac1 cyfip2-GFP, cyfip2^-/-^;Tg(∆Rac1 cyfip2-GFP), cyfip2^+/-^;Tg(∆FMRP cyfip2-GFP, cyfip2^-/-^;Tg(∆FMRP cyfip2-GFP), and fmr1^-/-^* after either no heatshock, or a 38°C heatshock at 30 hpf. G-J) Percent of non-responsive larvae in Figures 5A-D. Data are presented as mean ± SD. Comparisons made using Kruskal-Wallis test with Dunn’s multiple comparisons correction. * p < 0.05, **p < 0.01, ***p < 0.001, ****p < 0.0001.

**Supplementary Table S1. Genotyping**

| Allele | GeneID: | Primer Sequence | Annealing temp(°C) | Extension time (s) | Restriction Enzyme | Expected amplicon size (bp) |
| --- | --- | --- | --- | --- | --- | --- |
| cyfip2^p400^ | NCBI: 100002872  ZFIN: ZDB-GENE-080724-2 | Forward:  5 ′-CAAAGTCTTGCTGCGGATAAAAG-3′ | 52.8 | 30 | ApoI-HF (NEBL #R3566L) | WT: 149  MUT: 125 |
|  |  | Reverse:  5′-CTGCACCATCTGCTCACACAAATT-3′ |  |  |  |  |
| GFP | ZFIN: ZDB-EFG-070117-2 | Forward:  5 ′-GACGTAAACGGCCACAAGTT-3′ | 63 | 45 | n/a | 609 |
|  |  | Reverse:  5′- GAACTCCAGCAGGACCATGT-3′ |  |  |  |  |
| cyfip2^C179R^ | n/a | Forward:  5 ′-CTTCCAGCGCAAGGCCATTGAG-3′ | 53 | 30 | n/a | 293 |
|  |  | Reverse:  5′- GCAGACACTGTGTAATTCTGTTGTGATTGG-3′ |  |  |  |  |
| cyfip2^K723E^ | n/a | Forward:  5 ′-GCCCTCCATGATGGAATACGTGTTATATC-3′ | 52 | 30 | n/a | 431 |
|  |  | Reverse:  5′- GCCATTCAAGTTCCACTATGGAGGTAAG-3′ |  |  |  |  |
